# Cortical thickness, gray matter volume and cognitive performance: a cross-sectional study of the moderating effects of age on their inter-relationships

**DOI:** 10.1101/2022.09.29.510169

**Authors:** Marianne de Chastelaine, Sabina Srokova, Mingzhu Hou, Amber Kidwai, Seham S. Kafafi, Melanie L Racenstein, Michael D Rugg

## Abstract

In a sample comprising younger, middle-aged and older cognitively healthy adults (N = 375), we examined associations between mean cortical thickness, gray matter volume (GMV) and performance in four cognitive domains - memory, speed, fluency and crystallized intelligence. In almost all cases, the associations were moderated significantly by age, with the strongest associations in the older age group. An exception to this pattern was identified in a younger adult sub-group aged less than 23 yrs, when a negative association between cognitive performance and cortical thickness was identified. Other than for speed, all associations between structural metrics and performance in specific cognitive domains were fully mediated by mean cognitive ability. Cortical thickness and GMV explained unique fractions of the variance in mean cognitive ability, speed and fluency. In no case, however, did the amount of variance jointly explained by the two metrics exceed 7% of the total variance. These findings suggest that cortical thickness and GMV are distinct correlates of domain-general cognitive ability, that the strength and, for cortical thickness, the direction of these associations is moderated by age, and that these structural metrics offer only limited insights into the determinants of individual differences in cognitive performance across the adult lifespan.

---

The effects of age on brain structure, and their role in mediating age-related cognitive decline, has long been a topic of interest. With the development of semi-automated pipelines for the analysis of structural MRI data (Fischl and Dale, 2000) much of this interest has focused on the structural metric of neocortical (henceforth cortical) thickness. Consequentially, a large literature has developed documenting the trajectory of cortical thickness across the lifespan. As summarized in a report that aggregated data across more than 100 cross-sectional and longitudinal datasets, cortical thickness peaks in late infancy and declines monotonically with increasing age thereafter (Bethlehem et al., 2022).

In contrast to the lifelong negative association between cortical thickness and age evident after early childhood, the association between cortical thickness and cognitive ability across the lifespan has been reported to be non-linear. Convergent findings from cross-sectional and longitudinal studies indicate that cortical thickness is negatively correlated with cognitive ability in adolescence and young adulthood (de Chastelaine et al., 2019; Krogstrud et al., 2021; Schnack et al., 2015; Tamnes et al., 2010) but demonstrates a positive correlation with cognitive ability in later life (e.g., de Chastelaine et al., 2019; Karama et al., 2014; Salthouse et al., 2015; Sun et al., 2016). The findings for young individuals have been interpreted as a reflection of the beneficial effects of relatively extensive synaptic ‘pruning’ during cortical development and maturation (Schnack et al., 2015; Tadayon et al., 2020). The findings for older adults are consistent with the intuition that ‘more is better’ and the widely held view that degradation of brain structure is a mediator of age-related cognitive decline (the corollary of the ‘brain maintenance’ hypothesis, Cabeza et al., 2018). The notion of cortical thinning as a mediator of individual differences in age-related cognitive decline has however been called into question by longitudinal evidence that childhood intelligence accounts for most of the association between intelligence and cortical thickness some 60 years later, suggesting a lifelong association between the two variables (Karama et al., 2014).

The studies discussed above employed relatively undifferentiated metrics of thickness, such as the mean thickness of the entire cortical mantle. Numerous studies have examined associations at a more granular level, reporting correlations between the thickness of specific cortical regions and performance in different cognitive domains (e.g., Burzynska et al., 2012; Vonk et al., 2019). However, the reliability of such correlations has been called into question (Masouleh et al., 2022). Moreover, many studies reporting regionally specific associations between thickness and cognitive performance did not account for the strong correlations that exist between regional thickness measures. When this inter-dependency is accounted for, little evidence of regionally specific associations between cortical thickness and cognitive performance remains (Hou et al., 2021; Kranz et al., 2018; Krogsrud et al., 2021; Salthouse et al., 2015; but see Lee et al., 2016). Analogously, associations between thickness and specific cognitive abilities are also largely eliminated after the variance shared across different cognitive measures is controlled for (Hou et al., 2021; Krogsrud et al., 2021; Tsapanou et al., 2019; Salthouse et al., 2015). Thus, findings indicative of selective associations between regional cortical thickness and domain-specific cognitive performance seem largely to reflect a single association between global cortical thickness and mean cognitive ability.

The employment of cortical thickness as a structural brain metric is a relatively recent development. A variety of measures related to brain size or volume have a longer, and somewhat independent, history as predictors of cognitive ability, mainly the prediction of general intelligence. Volumetric measures decline monotonically with increasing age from mid-childhood (Bethlehem et al., 2022), and they have consistently been reported to correlate with cognitive ability seemingly independently of sex or age (for reviews and meta-analyses, see Pietschnig et al., 2015, 2022). Findings from one recent large-scale study (Nave et al., 2018) indicated that the principal driver of the relationship between brain volume and IQ is the volume of gray matter (GMV). As with cortical thickness, associations between brain volume and cognition at the regional level appear to be relatively weak, if reliable at all (Cox et al., 2019; Masouleh et al., 2019), and largely reduce to an association captured by whole brain measures.

Here, we took advantage of structural MRI and neuropsychological test data obtained from 375 participants, falling mainly into the younger (18-30 yrs) and older adult (ca. 65-75 yrs) age ranges, along with a smaller middle-aged group (45-55 yrs), to examine inter-relationships between cortical thickness, GMV and cognitive performance. As noted above, there is scant evidence that measures of cortical thickness or GMV demonstrate reliable associations with cognitive ability at the regional level. Accordingly, we focus here on whole brain metrics of these variables. We address four principal questions. First, with the additional power afforded by a larger sample, do we replicate our prior findings (de Chastelaine et al., 2019) that the relationship between cortical thickness and cognitive ability is moderated by age, such that a negative correlation is found in young adulthood while a positive relationship is evident in later life? Second, can we replicate prior findings (Hou et al., 2021; Salthouse et al., 2015) that cortical thickness is associated with general rather than domain-specific aspects of cognitive ability, and do these findings extend to GMV? Third, is the relationship between GMV and cognitive ability moderated by age? Finally, do cortical thickness and GMV explain unique fractions of variance in cognitive performance, that is, are they distinct or redundant correlates of cognitive ability? To our knowledge, this last question has rarely been addressed (but see Hedden et al., 2016; Ritchie et al., 2015).

## Materials and Methods

### Participants

A total of 375 cognitively healthy adults contributed to the analyses presented below. The participants were recruited from UT Dallas and its surrounding communities, and comprised 195 younger adults (18-30 yrs), 145 older (63-76 yrs) adults, and 35 middle-aged (43-55 yrs) individuals. Demographic details (along with neuropsychological test scores – see below) are summarized in Table 1. Data from an additional 18 participants (5 younger, 3 middle-aged and 10 older) were rejected because of the low quality of their structural brain images.

**Table 1.** Demographic information and neuropsychological test scores (sum or mean(sd)) in young, middle-aged, and older adults.

|  | Young | Middle | Older |
| --- | --- | --- | --- |
| N | 195 | 35 | 145 |
| Sex (M/F) | 97/98 | 15/20 | 67/78 |
| Age at Neuropsychological test | 22.79 (3.13) | 49.60 (3.45) | 68.50 (3.44) |
| Years of Education | 15.71 (2.10) | 16.49 (2.48) | 16.84 (2.27) |
| MMSE | 29.30 (0.87) | 29.34 (0.77) | 29.21 (0.88) |
| CVLT Short Delay Free Recall | 12.89 (2.26) | 11.57 (2.38) | 10.69 (3.02) |
| CVLT Long Delay Free Recall | 13.33 (2.34) | 12.29 (2.43) | 11.27 (2.92) |
| CVLT Short Delay Cued Recall | 13.44 (2.17) | 13.14 (1.99) | 12.07 (2.60) |
| CVLT Long Delay Cued Recall | 13.73 (2.14) | 12.97 (1.95) | 12.18 (2.49) |
| CVLT Recall Composite | 53.39 (8.39) | 49.97 (8.14) | 46.13 (10.42) |
| CVLT Recognition Hits | 15.36 (1.02) | 15.29 (0.83) | 14.83 (1.34) |
| CVLT Recognition FAs | 0.83 (1.15) | 1.91 (2.16) | 2.25 (2.34) |
| WMS Logical Memory I | 30.67 (6.39) | 29.00 (6.25) | 28.53 (4.88) |
| WMS Logical Memory II | 28.15 (6.60) | 25.49 (5.90) | 25.63 (5.78) |
| WMS Composite | 58.82 (12.58) | 54.49 (11.76) | 54.17 (10.17) |
| F-A-S | 45.62 (11.47) | 47.11 (11.72) | 45.30 (11.71) |
| SDMT | 63.14 (11.26) | 55.17 (7.60) | 48.98 (8.40) |
| Trails A (Seconds) | 21.20 (7.29) | 23.97 (6.29) | 31.03 (10.85) |
| Trails B (Seconds) | 46.55 (18.30) | 51.89 (16.89) | 68.90 (26.66) |
| Forward/Backward Digit Span | 19.11 (4.22) | 17.97 (3.48) | 18.16 (3.87) |
| Category Fluency Test | 24.52 (5.33) | 23.60 (5.92) | 21.76 (5.19) |
| TOPF/WTAR | 108.60 (5.72) | 111.34 (4.95) | 111.05 (5.70) |
| Raven's | 11.08 (1.02) | 10.20 (1.68) | 9.33 (2.12) |

The participants were recruited into one or more of the fMRI studies conducted in our laboratory between 2011 and 2019. Participants were right-handed, had normal or corrected-to-normal vision, and were fluent English speakers before the age of five. Additionally, participants had no history of neurological or psychiatric disease, substance abuse, diabetes, or current or recent use of prescription medication affecting the central nervous system. The participants were accepted into a study according to a common set of inclusion and exclusion criteria that were intended to minimize the likelihood of including individuals with cognitive impairment arising from neuropathology, as determined by our neuropsychological test battery (see *Neuropsychological test battery* below).

The same MRI scanner and closely similar acquisition sequences were employed to acquire a T1-weighted structural MRI scan from each participant. For those participants who contributed to multiple studies, we employed the MRI scan that was acquired closest to the time they first undertook the neuropsychological test battery (mean time between the test and scan sessions was 4.0 weeks, range 0 – 43 weeks).

Structural brain data in the form of cortical thickness estimates have been reported previously for a sub-set of the present sample (36 of the younger participants, the 35 middle-aged participants included here, and 62 of the older participants; see de Chastelaine et al. 2019 and Hou et al., 2021). Neither the analyses of cortical thickness estimates from the full sample, nor the analyses of GMV, have been reported previously.

### Neuropsychological test battery

With the exception of one younger adult, participants completed the neuropsychological test battery on a separate day from the MRI session. The battery comprised the Mini-Mental State Examination (MMSE), the California Verbal Learning Test-II (CVLT; Delis et al., 2000), Wechsler Logical Memory (Tests 1 and 2; Wechsler, 2009), the Symbol Digit Modalities test (SDMT; Smith, 1982), the Trail Making (Test A and B; Reitan and Wolfson, 1985), the F-A-S subtest of the Neurosensory Center Comprehensive Evaluation for Aphasia (Spreen and Benton, 1977), the Wechsler Adult Intelligence Scale – Revised (WAIS; Forward and Backward digit span subtests; Wechsler, 1981), Category Fluency test (Benton, 1968), Raven’s Progressive Matrices (List 1; Raven et al., 2000). In addition, participants completed either the Wechsler Test of Adult Reading (WTAR, Wechsler, 2001) or its revised version, the Wechsler Test of Premorbid Functioning (TOPF; Wechsler, 2011). 348 participants completed the WTAR, and 27 completed the TOPF. The TOPF and WTAR scores were converted to commensurate measures by scaling the raw scores according to the two test’s respective age-scaled norms for age 43 years (the mean age of our participants). Scores on each of the neuropsychological tests are summarized in Table 1 for each age group. To minimize the likelihood of including participants with mild cognitive impairment or early dementia, potential participants were excluded prior to the MRI session if they performed > 1.5 SD below age norms on 2 or more non-memory tests, > 1.5 SD below the age norm on at least 1 memory test, or if their MMSE score was below 26. Because the four CVLT recall scores were highly correlated across participants (min r = 0.81), they were summed to give a single composite CVLT recall score prior to further analysis. For the same reason, the two WMS Logical Memory Scores (r = 0.86), were summed to give a single composite score. By contrast, the correlation between the two scores (A & B) comprising the Trails test was markedly lower (0.51); hence these scores were not averaged prior to entry in the PCA (see next section).

### Principal Components Analysis

The composite CVLT and Logical Memory recall scores, along with all other test scores apart from the MMSE, were subjected to principal components analysis (PCA) to identify the latent constructs underpinning performance on the test battery (cf. de Chastelaine et al., 2019; Hou et al., 2021). The MMSE was excluded from the PCA because of score compression, with more than 80% of the sample scoring at (30) or near (29) ceiling. The PCA was performed as follows: first, the test scores for each age group were z-transformed within-group (to obtain a solution that reflects relative test performance independently of age group). Second, the resulting z-scores were combined into a single matrix and subjected to a PCA as implemented in IBM SPSS Statistics v.28. Components were retained if their eigenvalues exceeded (or, for one component, closely approached) 1 (see Table 2). Third, the retained components were subjected to varimax rotation to simplify the solution space. In a final step, we computed participant-specific component scores to reflect the performance of a given participant on each of the retained principal components. To do this, we z-scored the raw test scores across age-groups and, for each of the retained components, multiplied the z-scored data by the component loading and summed the resulting scores.

**Table 2.** Varimax-rotated factor loadings from the PCA of the neuropsychological test battery.

|  | Memory | Speed | Fluency | Crystallized IQ |
| --- | --- | --- | --- | --- |
| CVLT Recall Composite | 0.85 | 0.11 | 0.24 | 0.01 |
| CVLT Recognition Hits | 0.70 | 0.01 | -0.10 | 0.33 |
| CVLT Recognition FAs | -0.65 | -0.11 | -0.24 | 0.28 |
| WMS Composite | 0.63 | 0.16 | 0.05 | 0.07 |
| F-A-S | -0.03 | 0.17 | 0.61 | 0.33 |
| SDMT | 0.18 | 0.72 | 0.28 | 0.06 |
| Trails A | -0.07 | -0.75 | -0.14 | 0.01 |
| Trails B | -0.13 | -0.73 | -0.04 | -0.28 |
| WAIS | 0.06 | 0.22 | 0.07 | 0.75 |
| Category Fluency | 0.11 | 0.21 | 0.76 | -0.08 |
| TOPF/WTAR | 0.07 | 0.01 | 0.46 | 0.64 |
| Raven's | 0.19 | 0.07 | 0.53 | 0.13 |
| Eigenvalues (% Variance) | 3.37 (28.08%) | 1.56 (13.02%) | 1.08 (9.01%) | 0.97 (8.10%) |

### MRI acquisition

A Philips Achieva 3T MR scanner (Philips Medical System, Andover, MA USA) equipped with a 32-channel head coil was used to acquire the anatomical images. T1-weighted anatomical images were acquired with a MPRAGE pulse sequence with the following range of parameters (due to incremental software upgrades, there were small differences in TR, TE and slice number across experiments): TR = 8.05-8.38 ms, TE = 3.68-3.90 ms, reconstructed FOV= 256×256, voxel size 1×1×1 mm, 160/176 slices, sagittal acquisition.

### Estimation of cortical thickness and brain volume

Cortical thickness and the different brain volume estimates were derived from each participant’s T1-weighted image using Freesurfer’s (v6.0) semi-automatic processing pipeline (http://surfer.nmr.mgh.harvard.edu/fswiki; Dale et al., 1999; Fischl and Dale, 2000; Fischl et al., 2002). After skull-stripping and intensity normalization each 3-dimensional T1 volume was reconstructed as an inflated 2-dimensional surface so that white matter and pial surfaces could be identified for each hemisphere. Using these surface estimates, cortical thickness could then be estimated.

After completion of this first step, white matter and pial surfaces were visually inspected by one of 6 trained raters who were blind to the participants’ age. Although the automated analysis generally succeeded in identifying white and gray matter boundaries, some brain regions (e.g., orbito-frontal, insula and temporal regions, and boundaries between gray matter and the pial surface and the cerebellum) frequently required further manual edits. Tissue classification errors were corrected by editing the white matter and brain masks returned by the first pass through the Freesurfer pipeline. In addition, ‘control points’ were added to voxels on, or close to, what were determined to be white matter paths that had been excluded by the automated analysis. Manual edits were repeated as necessary. Once completed, each edited scan was checked by a second rater for quality control purposes. Reliability of mean cortical thickness estimates across the 6 raters was assessed for 5 randomly selected brain images (two younger, two older and one middle-aged participant). The reliability coefficient (ICC[2]; Shrout and Fleiss, 1979) was 0.88.

Cortical thickness was measured as the distance from the grey/white matter boundary to the pial surface on a vertex-by-vertex basis across the entire cortical mantle of each hemisphere. Mean whole-brain cortical thickness was calculated by averaging cortical thickness values generated for the two hemispheres. GMV was indexed by the ‘Total Gray’ statistic returned by the Freesurfer package. This statistic is an aggregate of surface-based volume estimates (derived from the parcellated cortical ribbon including the hippocampus) and gray matter voxel counts from the segmented subcortical structures. We also extracted estimates of cortical, total brain and white matter volume. As reported in the Supplemental Materials, findings for cortical and total brain volume were somewhat weaker than, but very similar to, those for GMV, while the findings for white matter volume (WMV) were weaker still.

### Statistical analyses

Sex and age group differences in cognitive and neural metrics were evaluated with 3 (age-group) by 2 (sex) factorial ANOVAs. Significant age group effects were followed up as necessary with post-hoc t-tests. Relationships between brain metrics and performance in each of the four cognitive domains identified by PCA of the neuropsychological test scores (see below) were evaluated through a two-step hierarchical regression approach. The first step included age group, sex and years of education as predictor variables. The second step added the brain metric of interest (either thickness or GMV) and its interaction with age group as additional predictors. A significant increase in the variance explained by the model after the inclusion of these predictors indicated that the brain metric accounted for unique variance in performance in the cognitive domain in question. In cases where the interaction was non-significant the full model was re-run after dropping the term. If the interaction term was significant group-wise partial correlations between the brain and performance metrics, controlling for sex and chronological age, were computed. The same approach was taken for the ‘mean cognitive ability’ metric formed by summing participants’ four component scores (see below).

The Bonferroni procedure was employed to correct the nominal significance level (p<.05) of the above-described regression and correlational analyses for multiple comparisons. The correction was applied separately for each brain metric (that is, the two metrics were treated as separate test families). For any follow-up group-wise correlations the correction was also applied separately for each age group. Because the five cognitive performance scores were highly correlated (rs between 0.378 and 0.905), we employed the ‘M_eff_ correction’ (Derringer, 2018; Nyholt, 2004) to estimate the Bonferroni divisor. This yielded a corrected significance level of p < 0.017.

To examine the specificity of any relationship between a brain metric and a given component score we re-ran the hierarchical regression analyses that identified the relationship after including the other 3 component scores as additional step 1 predictor variables (cf. Salthouse, 2015). The relationship was deemed specific if the addition of the brain metrics in step 2 of the analyses still led to a significant increase in explained variance.

## Results

### Neuropsychological test scores

Performance on the neuropsychological test battery is reported in Table 1.

### Principal Components Analysis

The varimax-rotated PCA solution returned 4 components that, between them, accounted for 58.21% of the variance in the neuropsychological test scores. The loadings of each test on the 4 rotated components, as well as their eigenvalues and proportion of variance explained, are reported in Table 2. It can be seen from the table that the components roughly correspond to the constructs of ‘memory’, ‘speed’, ‘fluency’ and ‘crystallized intelligence’ (a closely similar pattern of loadings was reported by de Chastelaine et al., 2019 following PCA of a sub-set of the present data). As in Hou et al. (2021), we generated a mean cognitive ability score for each participant by summing the four component scores. We also estimated general cognitive ability by using the loadings associated with the first unrotated principal component extracted from the PCA of the neuropsychological test data (cf. Nave et al., 2019). Controlling for sex, age group, and years of education, the two estimates of mean ability were almost perfectly correlated (r = 0.997). Hence, below, we only report the findings for the first of these metrics. We interpret it as a rough proxy for general intelligence.

### Outlier identification

We conservatively defined outlying data points as +/- >3.5 SDs from the mean for each age group. This criterion identified one younger participant with excessively low memory, fluency, and mean cognitive ability scores, and one older participant with an excessively low speed component score. No other behavioral or brain measures met the criterion. The outlying scores were dropped from all analyses reported below. Thus, analyses were conducted on the full sample (N = 375) in the case of the crystallized IQ component scores, and on a sample size of N = 374 for the remaining 3 component scores and mean cognitive ability.

### Cognitive component scores

Component scores are summarized by age group and sex in Table 3. Also summarized in the table are the mean cognitive ability scores (see above). Fixed effects ANOVAs were employed to examine the effects of age group (younger, middle aged and older) and sex on each component. The outcomes of the ANOVAs are summarized in Table 4, along with the results of post-hoc pairwise contrasts when these were indicated. As is evident from the table, memory scores varied robustly and additively according to the factors of sex and age group, such that females out-performed males, and there was a graded effect of age group, with younger adults demonstrating the highest scores and older adults the lowest. By contrast, for the speed and fluency contrasts, only a graded age group effect was evident. For crystallized IQ, the age group factor was non-significant, but a small sex effect was evident, with males demonstrating the higher scores. Finally, as in the case of speed and fluency, mean cognitive ability varied with age group but not sex. In light of the sex effects that were evident for two of the five cognitive constructs, we included sex as an independent variable in all of the regression and correlational analyses described below.

**Table 3.**
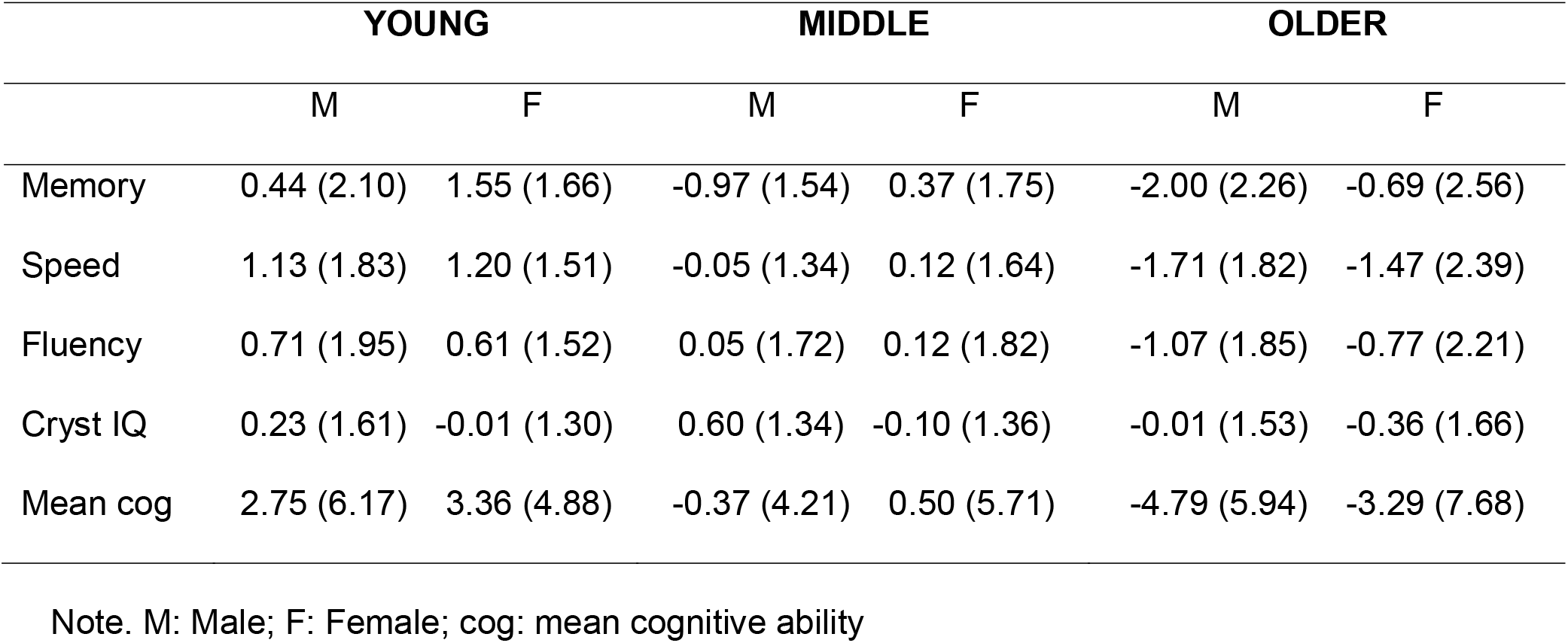
Component and mean cognitive ability scores (mean (sd)) stratified by age group and sex.

**Table 4.**
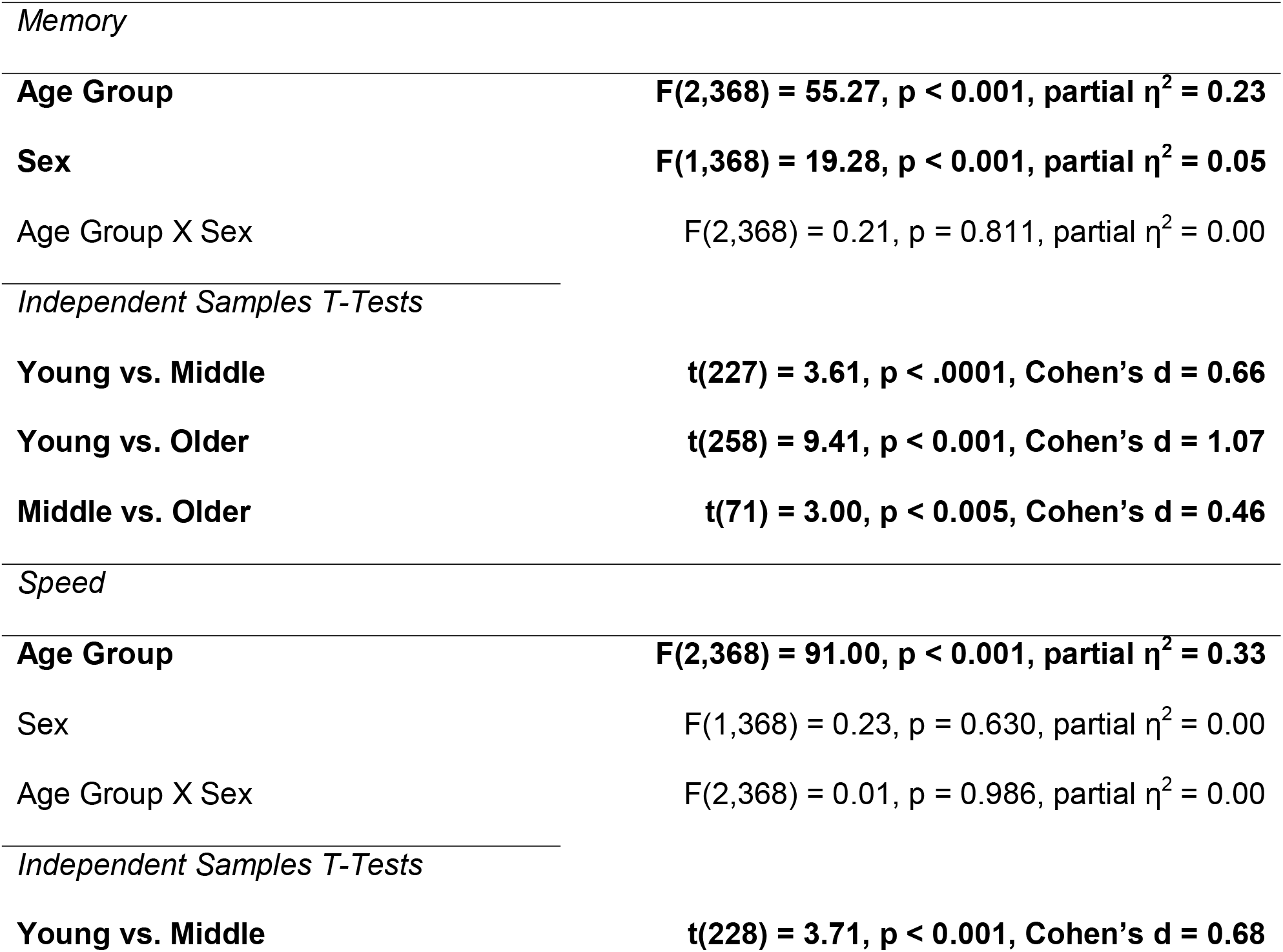
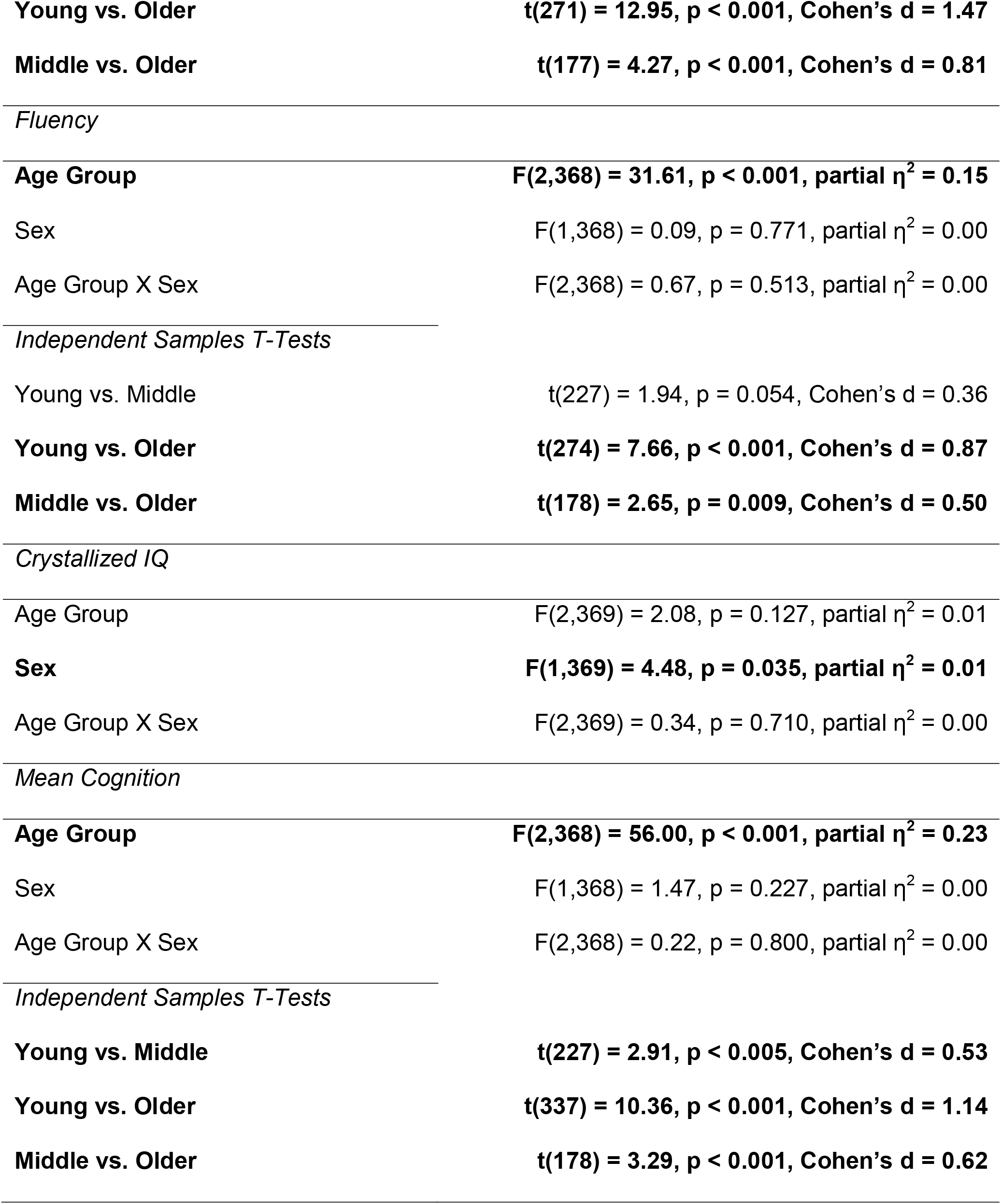
Results of fixed effects ANOVAs examining the effects of age group (young, middle aged and older) and sex on each cognitive component score, along with the outcomes of post-hoc pairwise contrasts (bold rows denote significance at p < 0.05).

### Years of education

ANOVA (factors of age group and sex) gave rise solely to a significant effect of age group [F(2,369) = 12.84, p < 0.001, partial η^2^ = 0.07]. Post-hoc tests revealed that years of education were significantly greater for the older than the younger group, with the middle-aged adults differing significantly from neither of the other groups. These findings motivated the inclusion of years of education as a control variable in the across-group regression and correlational analyses. However, all findings reported below were unaltered when this variable was omitted from the analyses.

### Cortical thickness

Whole-brain cortical thickness estimates are summarized in Table 5A and illustrated in Figure 1A. ANOVA of these data (factors of age group and sex) gave rise to a significant effect for age group [F(2,369) = 198.20, p < 0.001, partial η^2^ = 0.52], but no evidence of a sex effect (p = 0.402) or a sex x age group interaction (p = 0.167). Post-hoc contrasts revealed that thickness was graded across the age groups, such that the estimates for younger adults exceeded those for the middle-aged group [t(228) = 6.77, p < 0.001, Cohen’s d = 1.24] while, in turn, the middle-aged group demonstrated larger estimates than did the older age group [t(178) = 4.98, p < 0.001, Cohen’s d = 0.94].

**Table 5.** Cortical thickness (A) and gray matter volume (B) measures (mean (sd)) stratified by sex and age group.

|  | Young |  | Middle |  | Older |  |
| --- | --- | --- | --- | --- | --- | --- |
|  | Male | Female | Male | Female | Male | Female |
| A |  |  |  |  |  |  |
| Cortical Thickness | 2.54 | 2.53 | 2.42 | 2.44 | 2.32 | 2.34 |
| (mm) | (0.09) | (0.08) | (0.10) | (0.09) | (0.11) | (0.10) |
| B |  |  |  |  |  |  |
| Grey Matter Volume | 710.63 | 634.37 | 648.37 | 606.69 | 612.95 | 555.48 |
| (ml) | (56.55) | (49.13) | (47.50) | (36.55) | (43.85) | (49.32) |

**Figure 1.**
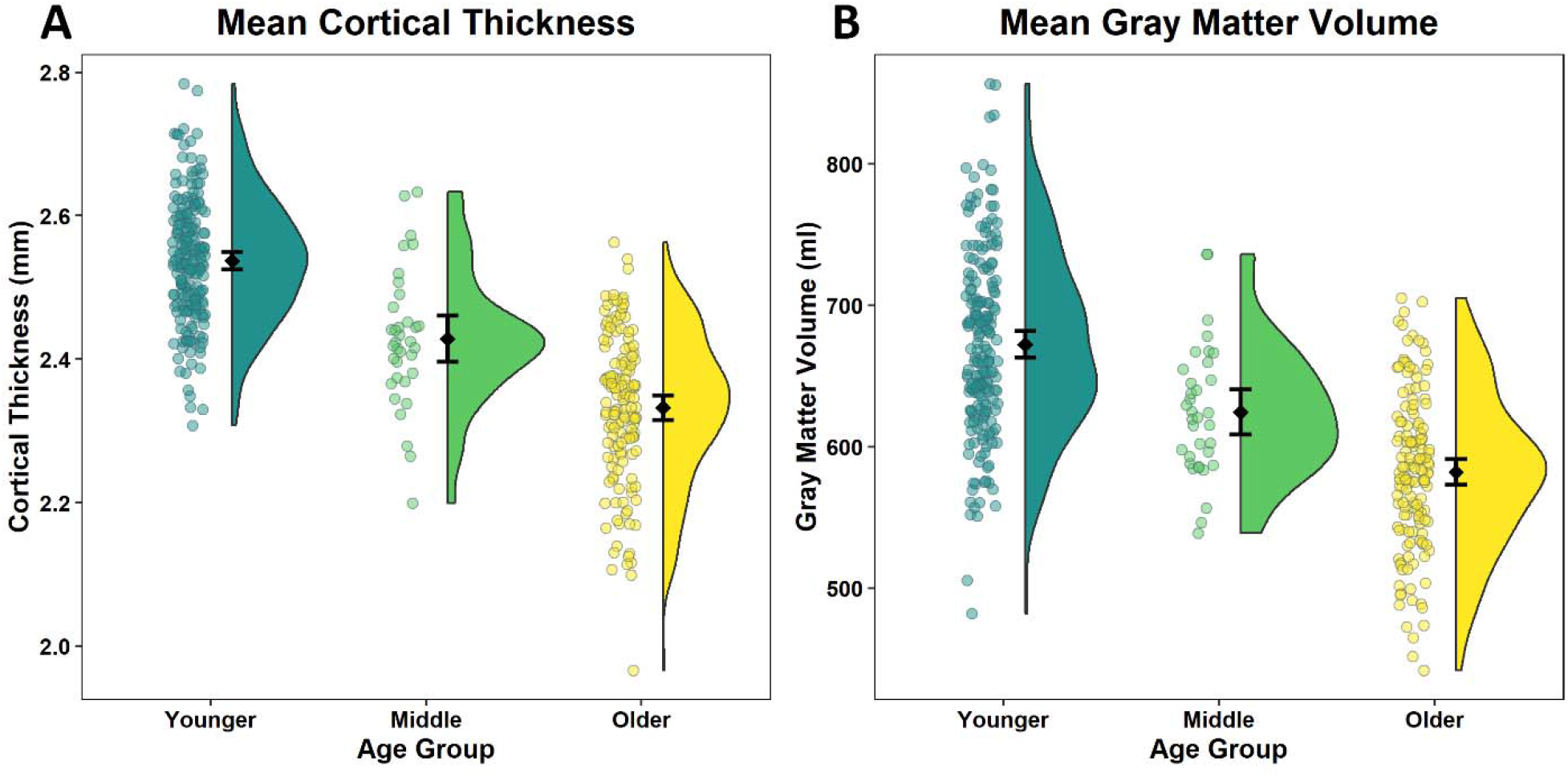
Distributions of estimates of cortical thickness (A) and GMV (B) in each age group. Black dots and bars represent means +/-95% confidence intervals.

### Gray matter volume

Estimates for GMV are summarized in Table 5B and illustrated in Figure 1B. ANOVA of GMV gave rise to significant effects of age group [F(2,369) = 130.98, p < 0.001, partial η^2^ = 0.42] and sex [F(1,369) = 75.46, p < 0.001, partial η^2^ = 0.17], with no evidence of an interaction between the factors (p = 0.076). As for cortical thickness, post-hoc contrasts indicated that GMV was lower in the middle-aged than the younger group [t(228) = 4.15, p < 0.001, Cohen’s d = 0.76], and lower in the older than the middle-aged group [t(178) = 4.24, p < 0.001, Cohen’s d = 0.80]. The main effect of sex was driven by greater GMV in males.

### Relationships between chronological age, cognitive performance and structural metrics

We used partial correlations (controlling for sex and education) to examine relationships between chronological age and cognitive and brain structural metrics separately for each age group. The results of these analyses are summarized in Table 6, where it is evident that in the younger and older age groups cortical thickness was significantly and negatively correlated with age (albeit more strongly in the older group; difference between correlation coefficients p < 0.05). By contrast, a reliable negative correlation between age and GMV is present in the older group only (while non-significant, a modest negative correlation is also evident in the middle-aged group). Turning to the cognitive component scores, no score demonstrated a reliable correlation with age in the younger age group. By contrast, except for crystallized IQ, all cognitive scores correlated reliably with age in the older group. Only in the case of the speed component, however, did the corresponding correlations differ significantly between the younger and older age groups (p < 0.05). Again, although they were non-significant, the pattern of correlations in the middle-aged group roughly mirrored that in the older group.

**Table 6.** Correlations between chronological age, cognitive performance and structural metrics for each age group, after controlling for sex and years of education (p values in parentheses).

|  | Chronological age |  |  |
| --- | --- | --- | --- |
|  | Young | Middle | Older |
| Mean Cortical Thickness | **-0.20 (0.007)** | -0.13 (0.468) | **-0.41 (< 0.001)** |
| GMV | -0.13 (0.082) | -0.20 (0.264) | **-0.36 (< 0.001)** |
| Memory | -0.04 (0.594) | -0.19 (0.296) | **-0.22 (0.009)** |
| Speed | -0.12 (0.111) | -0.16 (0.359) | **-0.34 (< 0.001)** |
| Fluency | -0.07 (0.306) | -0.16 (0.377) | **-0.20 (0.017)** |
| Crystallized IQ | -0.03 (0.634) | 0.05 (0.783) | -0.06 (0.458) |
| Mean Cognitive Ability | -0.08 (0.292) | -0.15 (0.391) | **-0.25 (0.002)** |

### Relationships between cortical thickness and cognitive performance

The outcomes of the hierarchical regression analyses predicting each cognitive score from mean cortical thickness and its interaction with age group are summarized in Table 7. As is evident from the table, sex, age group and years of education were each significant predictors of memory performance and mean cognitive ability, while age group and years of education predicted speed, fluency and crystallized IQ. Crucially, the inclusion of the thickness and thickness by age interaction terms in the model led to a significant increase in explained variance for each of the component scores and for mean cognitive ability (all ps < 0.005 other than for crystallized IQ, for which p = 0.018). Moreover, again in every case, the age group by thickness interaction term was significant.

**Table 7.**
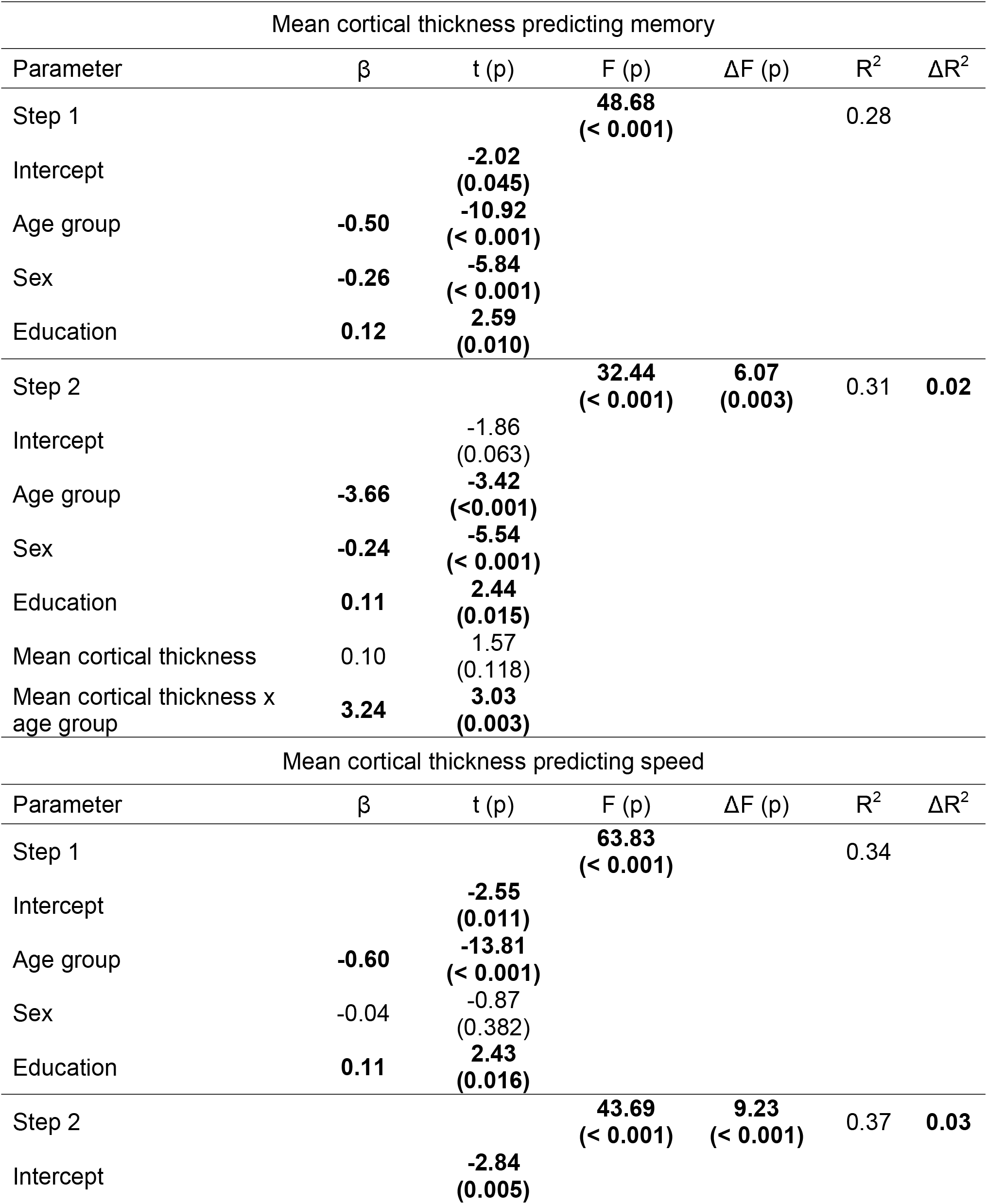
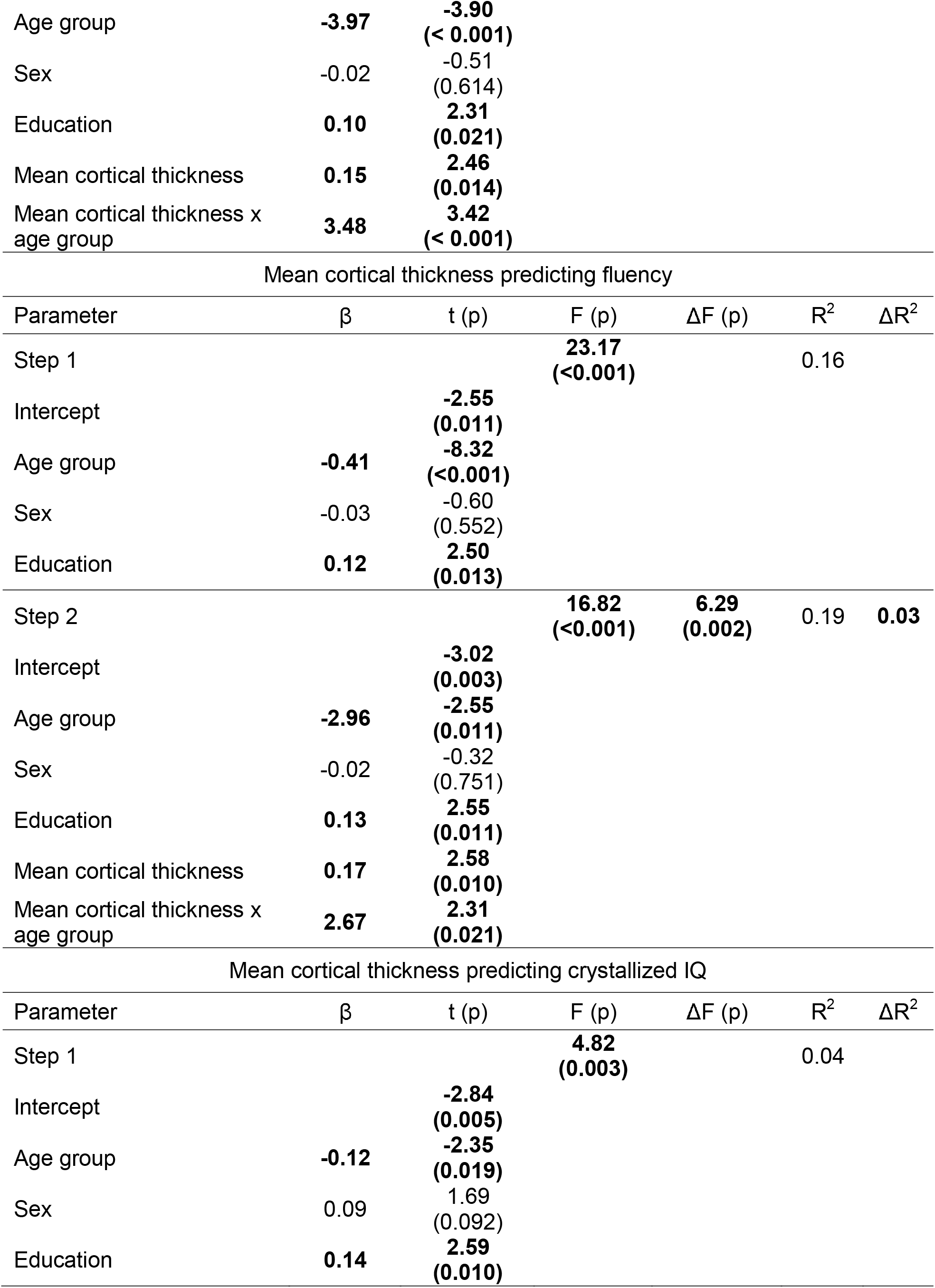
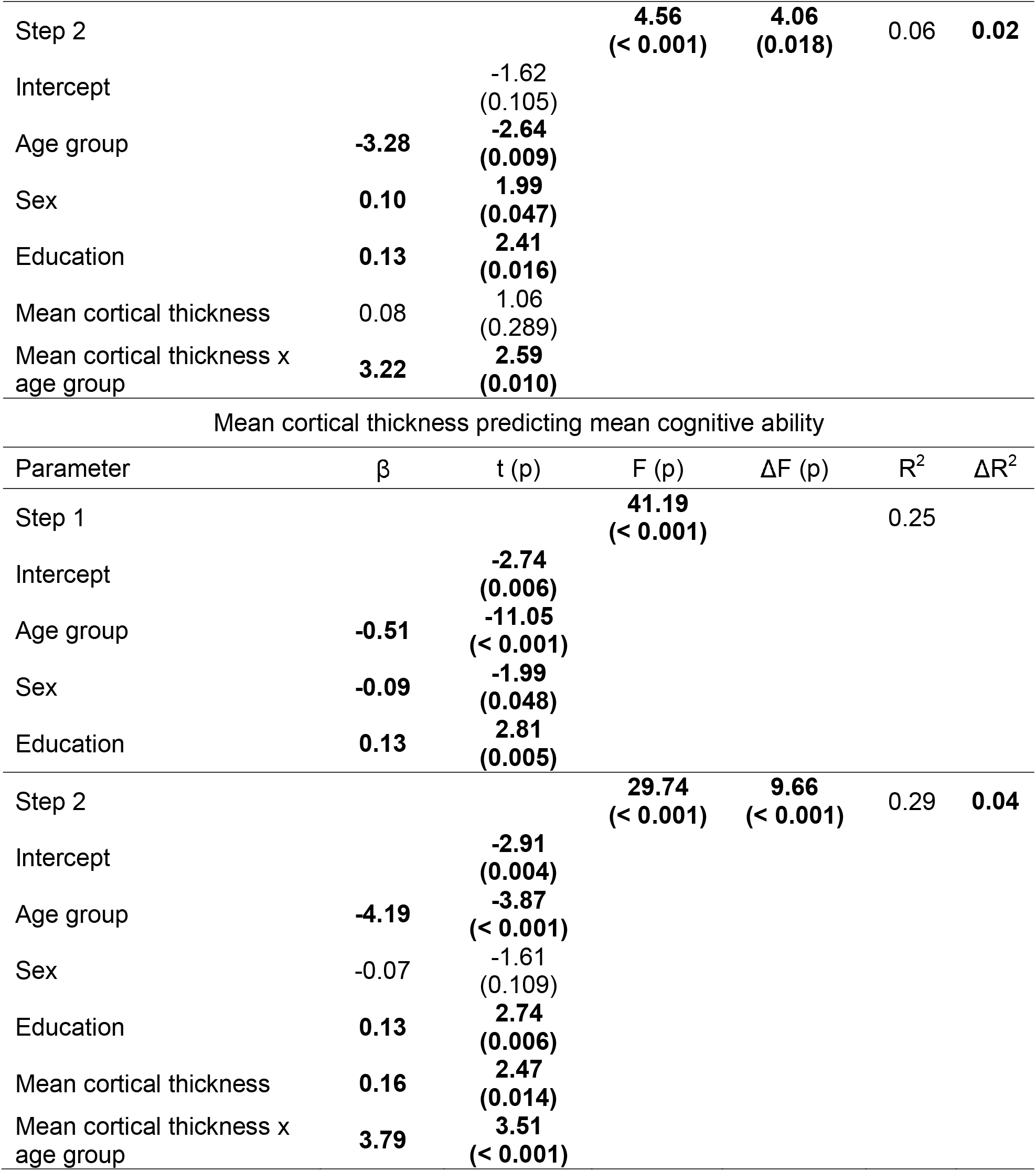
Hierarchical linear regression results for mean cortical thickness predicting cognitive performance (p values in parentheses).

| Mean cortical thickness predicting memory |  |  |  |  |  |  |
| --- | --- | --- | --- | --- | --- | --- |
| Parameter | $\beta$ | t (p) | F (p) | $\Delta F$ (p) | R<sup>2</sup> | $\Delta R^2$ |
| Step 1 |  |  | **48.68**<br>(**< 0.001**) |  | 0.28 |  |
| Intercept |  | **-2.02**<br>(**0.045**) |  |  |  |  |
| Age group | **-0.50** | **-10.92**<br>(**< 0.001**) |  |  |  |  |
| Sex | **-0.26** | **-5.84**<br>(**< 0.001**) |  |  |  |  |
| Education | **0.12** | **2.59**<br>(**0.010**) |  |  |  |  |
| Step 2 |  |  | **32.44**<br>(**< 0.001**) | **6.07**<br>(**0.003**) | 0.31 | **0.02** |
| Intercept |  | -1.86<br>(0.063) |  |  |  |  |
| Age group | **-3.66** | **-3.42**<br>(**<0.001**) |  |  |  |  |
| Sex | **-0.24** | **-5.54**<br>(**< 0.001**) |  |  |  |  |
| Education | **0.11** | **2.44**<br>(**0.015**) |  |  |  |  |
| Mean cortical thickness | 0.10 | 1.57<br>(0.118) |  |  |  |  |
| Mean cortical thickness x age group | **3.24** | **3.03**<br>(**0.003**) |  |  |  |  |
| Mean cortical thickness predicting speed |  |  |  |  |  |  |
| Parameter | $\beta$ | t (p) | F (p) | $\Delta F$ (p) | R<sup>2</sup> | $\Delta R^2$ |
| Step 1 |  |  | **63.83**<br>(**< 0.001**) |  | 0.34 |  |
| Intercept |  | **-2.55**<br>(**0.011**) |  |  |  |  |
| Age group | **-0.60** | **-13.81**<br>(**< 0.001**) |  |  |  |  |
| Sex | -0.04 | -0.87<br>(0.382) |  |  |  |  |
| Education | **0.11** | **2.43**<br>(**0.016**) |  |  |  |  |
| Step 2 |  |  | **43.69**<br>(**< 0.001**) | **9.23**<br>(**< 0.001**) | 0.37 | **0.03** |
| Intercept |  | **-2.84**<br>(**0.005**) |  |  |  |  |

|  |  |  |
| --- | --- | --- |
| Age group | **-3.97** | **-3.90**<br>**(< 0.001)** |
| Sex | -0.02 | -0.51<br>(0.614) |
| Education | **0.10** | **2.31**<br>**(0.021)** |
| Mean cortical thickness | **0.15** | **2.46**<br>**(0.014)** |
| Mean cortical thickness x<br>age group | **3.48** | **3.42**<br>**(< 0.001)** |

Mean cortical thickness predicting fluency
| Parameter | $\beta$ | t (p) | F (p) | $\Delta F$ (p) | R<sup>2</sup> | $\Delta R^2$ |
| --- | --- | --- | --- | --- | --- | --- |
| Step 1 |  |  | **23.17**<br>**(<0.001)** |  | 0.16 |  |
| Intercept |  | **-2.55**<br>**(0.011)** |  |  |  |  |
| Age group | **-0.41** | **-8.32**<br>**(<0.001)** |  |  |  |  |
| Sex | -0.03 | -0.60<br>(0.552) |  |  |  |  |
| Education | **0.12** | **2.50**<br>**(0.013)** |  |  |  |  |
| Step 2 |  |  | **16.82**<br>**(<0.001)** | **6.29**<br>**(0.002)** | 0.19 | **0.03** |
| Intercept |  | **-3.02**<br>**(0.003)** |  |  |  |  |
| Age group | **-2.96** | **-2.55**<br>**(0.011)** |  |  |  |  |
| Sex | -0.02 | -0.32<br>(0.751) |  |  |  |  |
| Education | **0.13** | **2.55**<br>**(0.011)** |  |  |  |  |
| Mean cortical thickness | **0.17** | **2.58**<br>**(0.010)** |  |  |  |  |
| Mean cortical thickness x<br>age group | **2.67** | **2.31**<br>**(0.021)** |  |  |  |  |

Mean cortical thickness predicting crystallized IQ
| Parameter | $\beta$ | t (p) | F (p) | $\Delta F$ (p) | R<sup>2</sup> | $\Delta R^2$ |
| --- | --- | --- | --- | --- | --- | --- |
| Step 1 |  |  | **4.82**<br>**(0.003)** |  | 0.04 |  |
| Intercept |  | **-2.84**<br>**(0.005)** |  |  |  |  |
| Age group | **-0.12** | **-2.35**<br>**(0.019)** |  |  |  |  |
| Sex | 0.09 | 1.69<br>(0.092) |  |  |  |  |
| Education | **0.14** | **2.59**<br>**(0.010)** |  |  |  |  |

| Step 2 |  |  | **4.56**<br>(**< 0.001**) | **4.06**<br>(**0.018**) | 0.06 | **0.02** |
| --- | --- | --- | --- | --- | --- | --- |
| Intercept |  | -1.62<br>(0.105) |  |  |  |  |
| Age group | **-3.28** | **-2.64**<br>(**0.009**) |  |  |  |  |
| Sex | **0.10** | **1.99**<br>(**0.047**) |  |  |  |  |
| Education | **0.13** | **2.41**<br>(**0.016**) |  |  |  |  |
| Mean cortical thickness | 0.08 | 1.06<br>(0.289) |  |  |  |  |
| Mean cortical thickness x<br>age group | **3.22** | **2.59**<br>(**0.010**) |  |  |  |  |
| Mean cortical thickness predicting mean cognitive ability |  |  |  |  |  |  |
| Parameter | $\beta$ | t (p) | F (p) | $\Delta F$ (p) | R<sup>2</sup> | $\Delta R^2$ |
| Step 1 |  |  | **41.19**<br>(**< 0.001**) |  | 0.25 |  |
| Intercept |  | **-2.74**<br>(**0.006**) |  |  |  |  |
| Age group | **-0.51** | **-11.05**<br>(**< 0.001**) |  |  |  |  |
| Sex | **-0.09** | **-1.99**<br>(**0.048**) |  |  |  |  |
| Education | **0.13** | **2.81**<br>(**0.005**) |  |  |  |  |
| Step 2 |  |  | **29.74**<br>(**< 0.001**) | **9.66**<br>(**< 0.001**) | 0.29 | **0.04** |
| Intercept |  | **-2.91**<br>(**0.004**) |  |  |  |  |
| Age group | **-4.19** | **-3.87**<br>(**< 0.001**) |  |  |  |  |
| Sex | -0.07 | -1.61<br>(0.109) |  |  |  |  |
| Education | **0.13** | **2.74**<br>(**0.006**) |  |  |  |  |
| Mean cortical thickness | **0.16** | **2.47**<br>(**0.014**) |  |  |  |  |
| Mean cortical thickness x<br>age group | **3.79** | **3.51**<br>(**< 0.001**) |  |  |  |  |

To elucidate the age group by thickness interactions we segregated the data by age group and, for each group, computed partial correlations between thickness and cognitive performance while controlling for sex, years of education, and chronological age. The correlations are given in Table 8A. As is evident from the table, significant correlations were identified for the older group only. The correlations were significant for all 5 cognitive scores before correction, and 2 correlations (crystallized IQ and mean cognitive ability) survived within-group FWE correction (p < 0.017). Scatterplots depicting these 2 correlations are illustrated in Figure 2.

**Table 8.** Correlations between cognitive performance and (A) cortical thickness and (B) GMV for each age group after controlling for chronological age, sex and years of education (p values in parentheses).

|  | A. Mean cortical thickness |  |  | B. GMV (adjusted for sex) |  |  |
| --- | --- | --- | --- | --- | --- | --- |
|  | Young | Middle | Older | Young | Middle | Older |
| Memory | -0.14<br>(0.062) | 0.09<br>(0.611) | **0.17**<br>**(0.041)** | -0.03<br>(0.723) | -0.14<br>(0.429) | 0.10<br>(0.249) |
| Speed | -0.09<br>(0.213) | 0.15<br>(0.421) | **0.19**<br>**(0.023)** | 0.12<br>(0.096) | 0.11<br>(0.557) | **0.26**<br>**(0.002)** |
| Fluency | -0.03<br>(0.635) | 0.21<br>(0.250) | **0.18**<br>**(0.035)** | 0.10<br>(0.177) | 0.04<br>(0.831) | **0.19**<br>**(0.021)** |
| Crystallized IQ | -0.07<br>(0.321) | -0.05<br>(0.783) | **0.20**<br>**(0.016)** | 0.03<br>(0.642) | -0.08<br>(0.660) | **0.22**<br>**(0.008)** |
| Mean Cognitive Ability | -0.10<br>(0.183) | 0.14<br>(0.457) | **0.23**<br>**(0.006)** | 0.06<br>(0.410) | -0.02<br>(0.907) | **0.22**<br>**(0.008)** |

**Figure 2.**
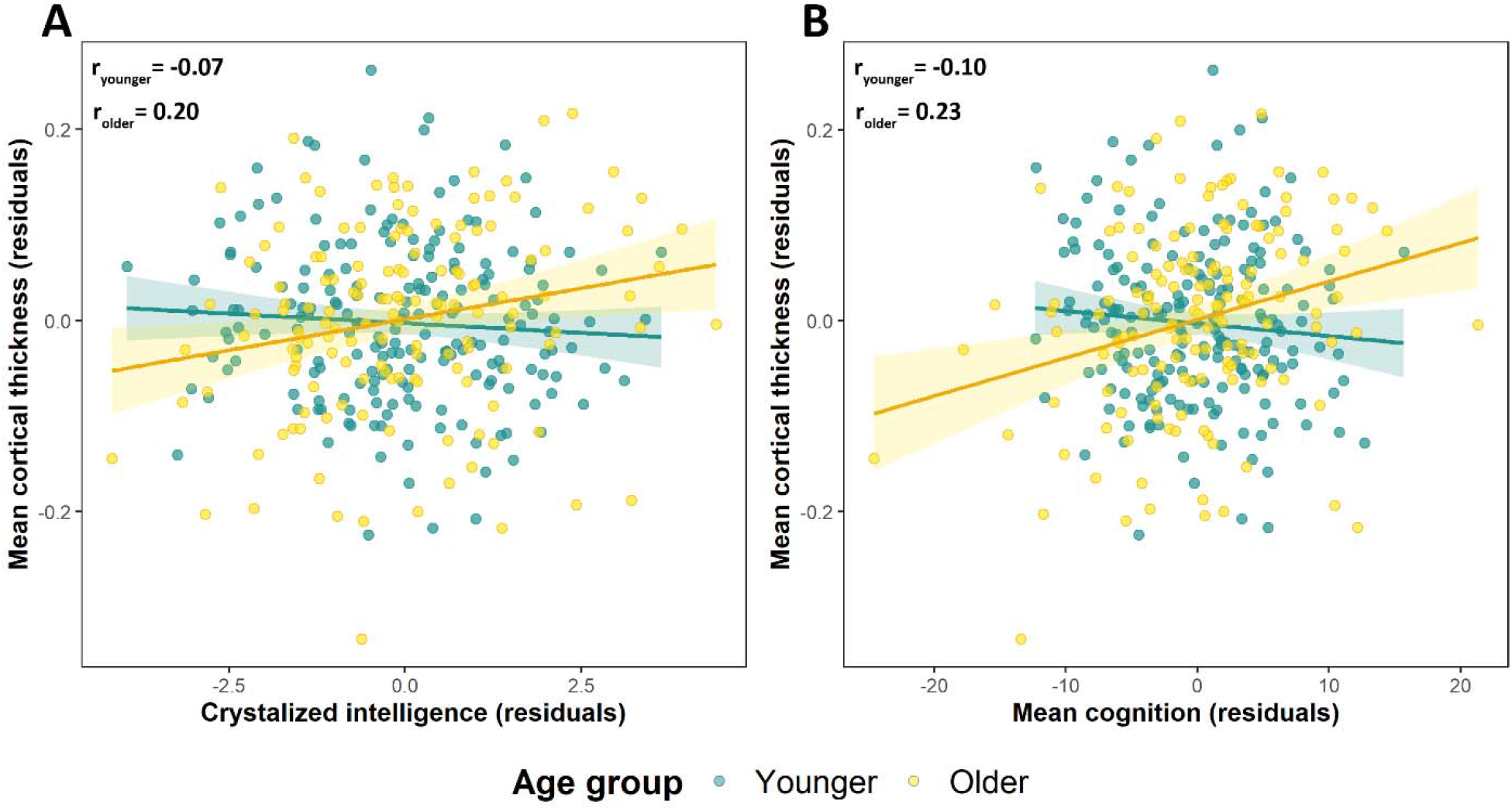
Scatter plots of the relationships between cortical thickness and component scores for crystallized intelligence (A) and mean cognitive ability (B) for younger and older age groups, controlling for chronological age, years of education and sex.

### Relationships between GMV and cognitive performance

As was reported above, GMV differed robustly according to sex. Accordingly, we adjusted GMV estimates for sex prior to submitting them to analyses. In so doing, we controlled for the variance in GMV that was directly attributable to sex while allowing variance attributable to other factors to remain. If this residual variance is shared with variance in cognitive performance, then the sex adjusted GMV metric will be identified as a significant predictor of performance (see Lee at al., 2019 for a closely analogous approach to controlling for sex differences in volumetric data).

The outcomes of the hierarchical regression analyses predicting cognitive performance from GMV and its interaction with age are summarized in Table 9. As can be seen from the table, GMV was a significant predictor of speed, fluency, and mean cognitive ability scores. In the case of fluency, the GMV x age group interaction was non-significant, and thus the model was re-run after dropping the term. The resulting partial correlation between GMV and fluency was highly significant (r369 = 0.17, p = 0.001) and a scatter plot of this relationship is illustrated in Figure 3A. By contrast, the interaction terms for speed and mean cognitive ability were both significant. The interactions were elucidated by group-wise partial correlations, which are shown in Table 8B. As is evident from the table, for each cognitive measure the correlations were largest and only reached significance in the older age group. The corresponding scatter plots are illustrated in Figures 3B and 3C.

**Table 9.**
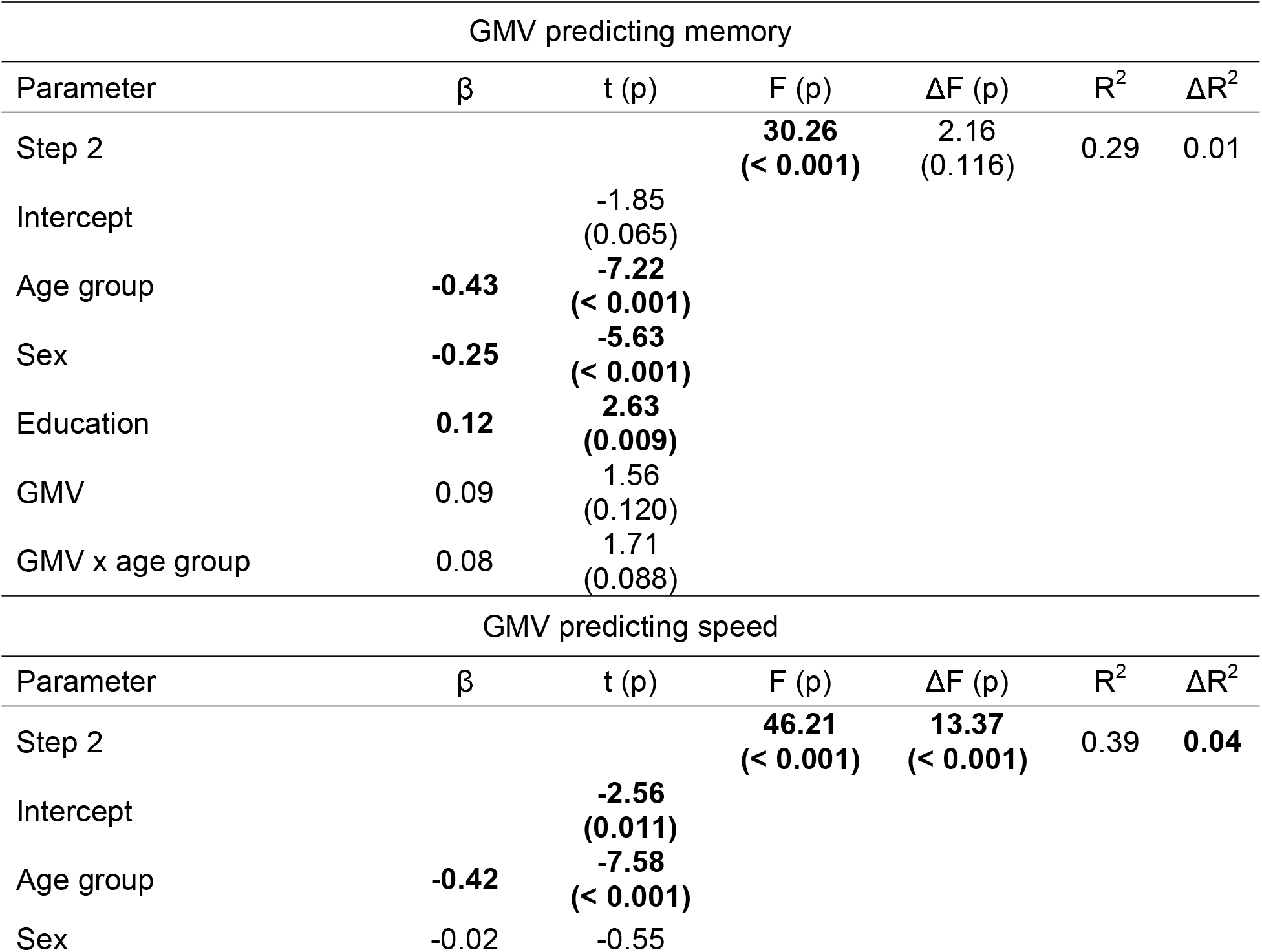
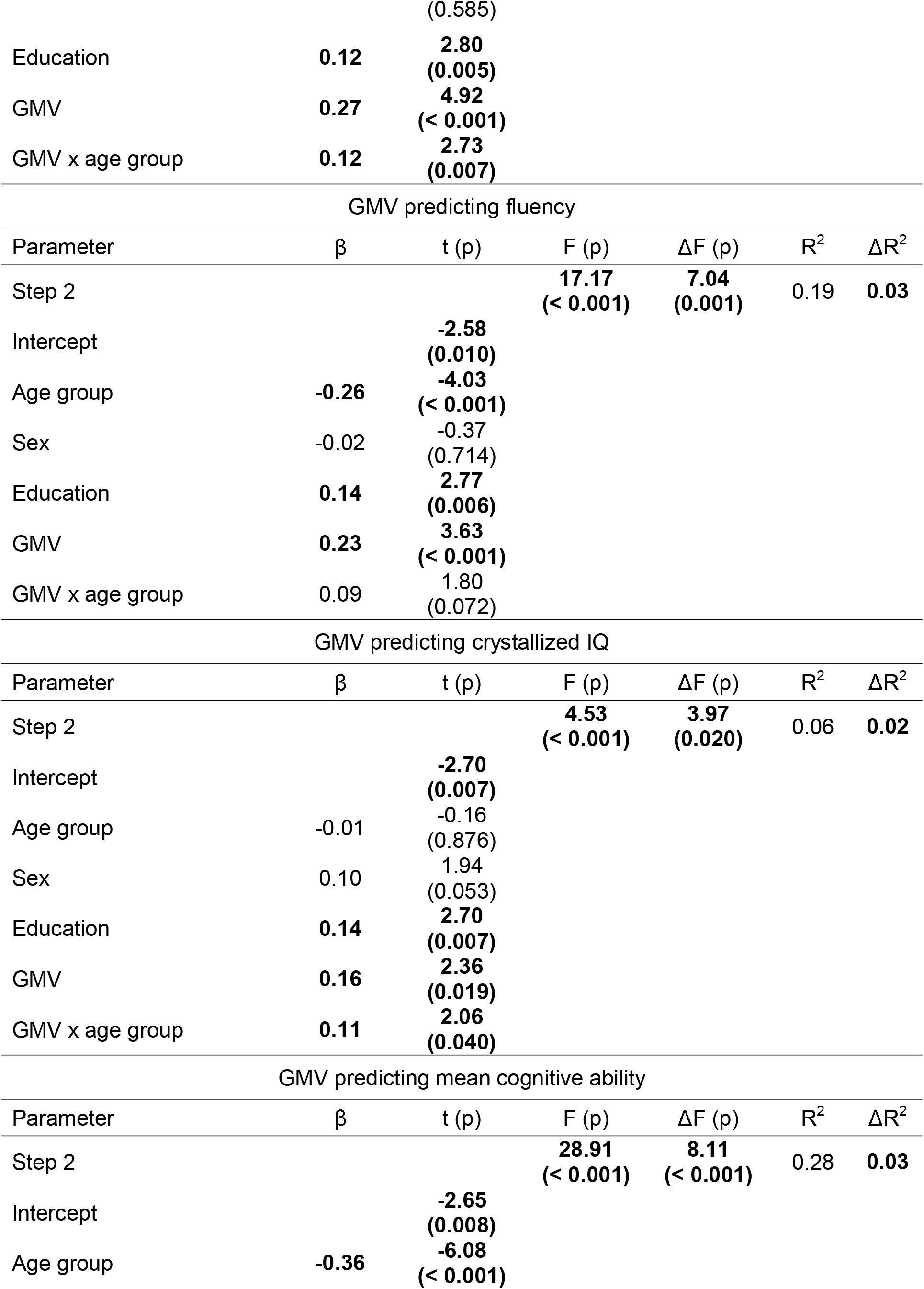
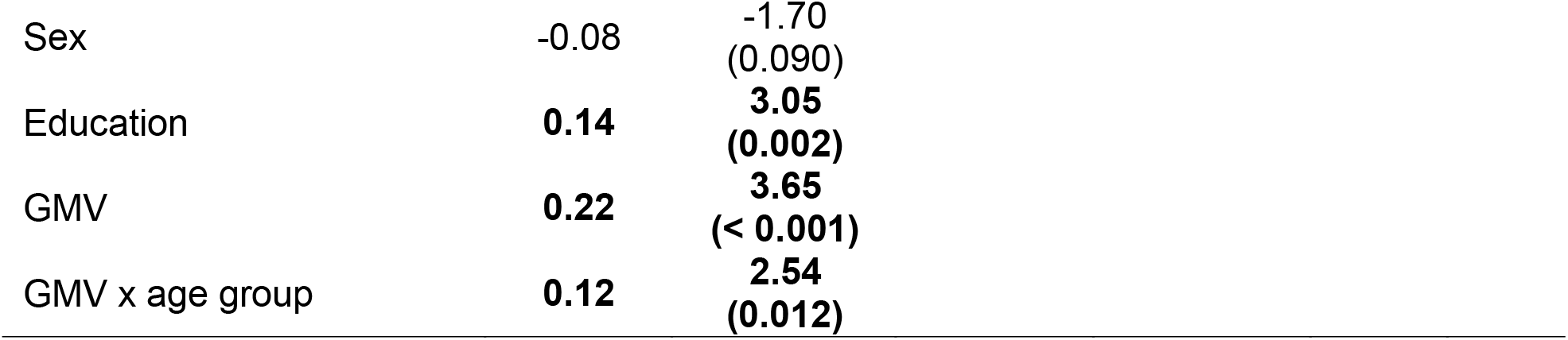
Hierarchical linear regression results for GMV (adjusted by sex) predicting cognitive performance (p values in parentheses, results of step 1 models are listed in Table 7).

| GMV predicting memory |  |  |  |  |  |  |
| --- | --- | --- | --- | --- | --- | --- |
| Parameter | $\beta$ | t (p) | F (p) | $\Delta F$ (p) | $R^2$ | $\Delta R^2$ |
| Step 2 |  |  | **30.26**<br>(**< 0.001**) | 2.16<br>(0.116) | 0.29 | 0.01 |
| Intercept |  | -1.85<br>(0.065) |  |  |  |  |
| Age group | **-0.43** | **-7.22**<br>(**< 0.001**) |  |  |  |  |
| Sex | **-0.25** | **-5.63**<br>(**< 0.001**) |  |  |  |  |
| Education | **0.12** | **2.63**<br>(**0.009**) |  |  |  |  |
| GMV | 0.09 | 1.56<br>(0.120) |  |  |  |  |
| GMV x age group | 0.08 | 1.71<br>(0.088) |  |  |  |  |
| GMV predicting speed |  |  |  |  |  |  |
| Parameter | $\beta$ | t (p) | F (p) | $\Delta F$ (p) | $R^2$ | $\Delta R^2$ |
| Step 2 |  |  | **46.21**<br>(**< 0.001**) | **13.37**<br>(**< 0.001**) | 0.39 | **0.04** |
| Intercept |  | **-2.56**<br>(**0.011**) |  |  |  |  |
| Age group | **-0.42** | **-7.58**<br>(**< 0.001**) |  |  |  |  |
| Sex | -0.02 | -0.55 |  |  |  |  |

|  |  | (0.585) |  |  |  |  |
| --- | --- | --- | --- | --- | --- | --- |
| Education | **0.12** | **2.80**<br>**(0.005)** |  |  |  |  |
| GMV | **0.27** | **4.92**<br>**(< 0.001)** |  |  |  |  |
| GMV x age group | **0.12** | **2.73**<br>**(0.007)** |  |  |  |  |
| GMV predicting fluency |  |  |  |  |  |  |
| Parameter | $\beta$ | t (p) | F (p) | $\Delta F$ (p) | R<sup>2</sup> | $\Delta R^2$ |
| Step 2 |  |  | **17.17**<br>**(< 0.001)** | **7.04**<br>**(0.001)** | 0.19 | **0.03** |
| Intercept |  | **-2.58**<br>**(0.010)** |  |  |  |  |
| Age group | **-0.26** | **-4.03**<br>**(< 0.001)** |  |  |  |  |
| Sex | -0.02 | -0.37<br>(0.714) |  |  |  |  |
| Education | **0.14** | **2.77**<br>**(0.006)** |  |  |  |  |
| GMV | **0.23** | **3.63**<br>**(< 0.001)** |  |  |  |  |
| GMV x age group | 0.09 | 1.80<br>(0.072) |  |  |  |  |
| GMV predicting crystallized IQ |  |  |  |  |  |  |
| Parameter | $\beta$ | t (p) | F (p) | $\Delta F$ (p) | R<sup>2</sup> | $\Delta R^2$ |
| Step 2 |  |  | **4.53**<br>**(< 0.001)** | **3.97**<br>**(0.020)** | 0.06 | **0.02** |
| Intercept |  | **-2.70**<br>**(0.007)** |  |  |  |  |
| Age group | -0.01 | -0.16<br>(0.876) |  |  |  |  |
| Sex | 0.10 | 1.94<br>(0.053) |  |  |  |  |
| Education | **0.14** | **2.70**<br>**(0.007)** |  |  |  |  |
| GMV | **0.16** | **2.36**<br>**(0.019)** |  |  |  |  |
| GMV x age group | **0.11** | **2.06**<br>**(0.040)** |  |  |  |  |
| GMV predicting mean cognitive ability |  |  |  |  |  |  |
| Parameter | $\beta$ | t (p) | F (p) | $\Delta F$ (p) | R<sup>2</sup> | $\Delta R^2$ |
| Step 2 |  |  | **28.91**<br>**(< 0.001)** | **8.11**<br>**(< 0.001)** | 0.28 | **0.03** |
| Intercept |  | **-2.65**<br>**(0.008)** |  |  |  |  |
| Age group | **-0.36** | **-6.08**<br>**(< 0.001)** |  |  |  |  |

|  |  |  |
| --- | --- | --- |
| Sex | -0.08 | -1.70<br>(0.090) |
| Education | **0.14** | **3.05**<br>**(0.002)** |
| GMV | **0.22** | **3.65**<br>**(< 0.001)** |
| GMV x age group | **0.12** | **2.54**<br>**(0.012)** |

**Figure 3.**
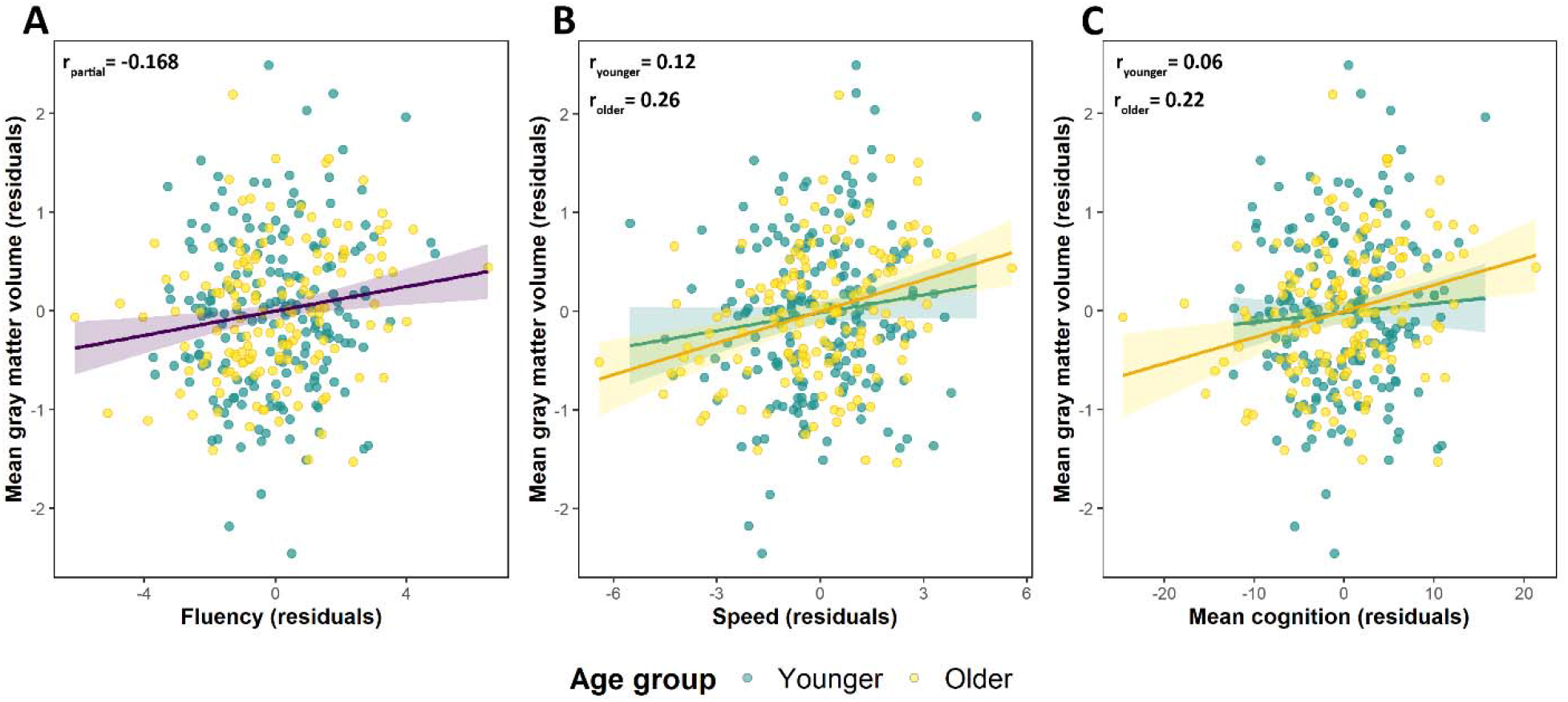
(A) scatter plot of the relationship across age groups between GMV and fluency component scores, controlling for age group, years of education and sex. (B&C): scatter plots of the relationships between GMV and the speed (B) and mean cognitive ability (C) component scores in the younger and older age groups, controlling for chronological age, years of education and sex.

### Specificity of the relationships between cognitive performance and brain metrics

As was discussed in the Introduction, it has previously been reported that the relationship between performance in a specific cognitive domain and structural brain metrics can be substantially attenuated when performance in other domains is controlled for. We addressed this issue here by re-running the hierarchical regression analyses predicting performance in each domain after including the other three component scores as additional step 1 predictors (see Methods). With the inclusion of these additional variables, cortical thickness and its interaction with age group no longer explained variance in any of the cognitive component scores (all ps > 0.336).

Similarly, after controlling for performance in the remaining cognitive domains, GMV no longer demonstrated an association with fluency (p = 0.090). By contrast, GMV continued to explain a significant (p = 0.001) fraction of the variance in the speed component scores.

### Cortical thickness and GMV as co-predictors of cognitive performance

Thickness and GMV were moderately correlated (controlling for sex and age group, r371 = 0.37, p < 0.001). Therefore, we examined whether the two metrics explained independent fractions of variance in the three cognitive scores – speed, fluency, and mean cognitive ability – with which they were independently associated. We constructed regression models in which cortical thickness, GMV, and their interactions with age group were employed to predict each component score (along with the control variables of age group, sex and years of education). In the initial iteration of these models the GMV by age interaction term did not approach significance and was therefore dropped. The GMV and the thickness x age group interaction terms were significant in the models predicting speed (both ps < 0.001), fluency (p = 0.014 and p = 0.022 for the GMV and thickness x age terms respectively) and mean cognitive ability (p = 0.021 and p < 0.001 for GMV and thickness x age group respectively). Thus, the two brain metrics accounted for a unique fraction of the variance in all three of the cognitive scores that were examined.

## Discussion

We examined the associations between mean cortical thickness, GMV and cognitive performance in relatively large samples of younger and older adults and a smaller sample of middle-aged individuals. In agreement with prior reports (see Introduction), age group moderated the relationship between cortical thickness and cognitive performance across each of the cognitive domains we examined. Also in agreement with prior findings, none of these relationships was reliable after controlling for variance in performance that was shared across domains. In the case of GMV, robust relationships with performance were evident for speed, fluency and mean cognitive ability, and two of these relationships were moderated by age group. In contrast with the findings for cortical thickness, in addition to demonstrating a robust correlation with mean cognitive ability, GMV explained unique variance in the speed cognitive component score. Finally, when entered as predictors into a common regression model, thickness and GMV explained independent fractions of variance in mean ability and in the speed and fluency component scores. Below, we discuss these findings in relation to the four questions that motivated the study.

Turning first to the findings for cortical thickness, we were able to confirm our prior findings (de Chastelaine et al., 2019) that positive associations with cognitive performance are evident in older but not in younger adults (we discuss the findings for the middle-aged sample in a separate section below). We also replicated previous reports (e.g., Hou et al., 2021; Tsapanou et al., 2019; Salthouse et al., 2015) that associations between cortical thickness and performance in different cognitive domains are largely mediated by a mean ability component reflecting variance shared across the different domains. However, with the enlarged younger adult sample available here (more than five times the size of our original sample) we failed to confirm our prior report of a negative association between thickness and cognitive ability in this age group.

A possible explanation for this null finding stems from the age distribution of our younger sample, which ranged from 18 to 30 years. According to Schnack et al. (2015), the negative association between cortical thickness and IQ peaks around age 20 years and declines over the following decade or so before reversing in direction in middle age. Given these findings, we split our younger adult sample at the median age (22 yrs) into younger (18-22 yrs; N = 101) and older (23-30 yrs; N = 94) sub-groups. The correlation (controlling for sex, chronological age and years of education) between mean cognitive ability (our proxy for IQ) and cortical thickness was small and non-significant in the older sub-group (r = 0.04), whereas the correlation was substantially larger and attained significance in the younger sub-group (r = −0.21, p = 0.036); the correlation with the thickness of the left hemisphere (the hemisphere for which Schnack et al. (2015) reported their strongest effects) was even larger: r = −0.28, p = 0.005. These findings, albeit from an unplanned and post-hoc analysis, are consistent with those of Schnack et al. in suggesting that the negative association between cortical thickness and cognitive performance dissipates during the third decade of life.

GMV demonstrated significant associations with cognitive performance in two cognitive domains (the exceptions being memory and crystallized IQ) and with mean cognitive ability. As indicated by reliable GMV by age group interactions terms, two of these associations (with speed and mean cognitive ability) were stronger in older than in younger adults. Indeed, we could find no evidence in our younger sample of a reliable correlation between GMV and any cognitive score. This included fluency, despite its interaction with age failing to attain significance in the regression model. Moreover, the correlations in the younger age group were, in the main, trivially small. These findings are at odds with the conclusions of two recent meta-analyses (Pietschnig et al., 2015, 2022), neither of which identified an effect of age on the association between brain volume and intelligence. The findings are not, however, the first to describe a moderating effect of age on the association between brain volume and cognitive ability.

Stronger associations in adults than children were reported in the meta-analysis of McDaniel (2005). And it was recently reported that, when combined with other structural metrics (although not independently), GMV was more strongly associated with intelligence in older than middle-aged adults (Cox et al., 2019).

It is unclear why we were unable to detect an association between cognitive performance and GMV in our younger adult sample given the prior evidence of a robust association in this age range. One possibility is that our inclusion and exclusion criteria resulted in range restriction at the lower end of the distributions of the cognitive performance scores of our younger age group. This possibility seems unlikely, however. Although the within-group variances of the speed and fluency scores were significantly greater in the older group (p = 0.027 and p = 0.05 respectively, according to Levene’s test), this was not the case for mean cognitive ability (p = 0.067).

Moreover, statistical evidence for truncation is lacking: according to Cohen’s (1959) formula, the ratio of the estimated unrestricted variance of the younger adults’ mean cognitive ability scores to the observed variance is 1.01. A second, related, possibility arises from the fact that most of the present younger participants were sampled from a college population, upwardly biasing cognitive performance relative to the general population (for example, performance on the age-normed tests in our neuropsychological battery averaged about 0.5 SD’s greater than the norms). We are unaware of any evidence pertaining to whether associations between brain volumetric measures and cognitive ability varies according to the average ability of the sample, let alone whether any such effect interacts with age. The present findings suggest however that examination of this question might prove worthwhile.

In contrast to the findings for cortical thickness, GMV remained a robust predictor of performance in a specific cognitive domain – speed – after controlling for the variance shared with the other domains. Of importance, the relationship between GMV and speed also remained significant (p = 0.002) - and was now unmoderated by age group (p = 0.196) - after controlling for mean cognitive ability. Thus, GMV demonstrated a robust association with speed independently of the contribution of the speed component to mean ability. This finding is especially remarkable given the high correlation between the speed and mean ability scores (r = 0.88, controlling for age group, sex and years of education).

The reasons for, and functional significance of, the association between speed and GMV are unclear, and we are unaware of any direct precedent in the literature (but see Brugulat-Serrat et al., 2019). In an effort to elucidate the association, we repeated the regression analysis described above using as independent variables the scores on the three tests - Trails A, Trails B and Symbol Digit Modalities - that loaded most heavily on the speed component. The results were unilluminating; each score demonstrated a statistically significant but small correlation with GMV that did not exceed the correlation obtained for the component score. Given prior evidence of an association between white matter integrity and processing speed (e.g., Madden et al., 2009; Kerchner et al., 2012), we also examined whether GMV was acting as a proxy for WMV (the two measures correlated at r = 0.65). As described in the Supplemental Materials, unlike GMV, which continued to be a robust predictor of speed scores, WMV did not explain any variance in speed when the two variables were entered as predictors into the same regression model. Therefore, it seems reasonable to conclude that it is GMV specifically that is a predictor of processing speed. The functional and neurobiological significance of this association are unclear.

Cortical thickness and GMV each accounted for significant fractions of the variance in cognitive performance, but the effect sizes were small (R^2^ ranging from around .03 to .04), as is typical in studies such as this (Pietschnig et al., 2022; Salthouse et al. 2015). The two metrics did however explain unique variance in three of the cognitive scores – speed, fluency, and mean ability – with which they demonstrated independent associations. Thus, cortical thickness and GMV appear to reflect distinct neurobiological correlates of cognitive performance. When thickness and GMV were employed as joint predictors of performance, the amount of explained variance remained modest - the largest fraction was for the speed component where, after controlling for age group, years of education and sex, the metrics jointly accounted for an additional 5.3% of the sample-wide variance. Within the older sample, for which correlations between the brain metrics and cognition were strongest, the variance in the speed component jointly explained by cortical thickness and GMV still only amounted to some 7% of the total after controlling for sex, age, and years of education. Therefore, it appears that these neural measures have the potential to provide only limited insight into the determinants of cognitive ability in cognitively healthy older individuals.

A major obstacle to understanding the significance of the associations between cognitive ability and cortical thickness and GMV identified here and in prior research is the paucity of knowledge about their neurobiological determinants. T1-weighted MRI provides no information about cortical microstructure, let alone the microstructural bases of individual differences in cortical thickness at different stages in the lifespan (the reversal in the direction of the relationship between thickness and cognitive ability in early relative to late adulthood strongly suggests that the determinants of thickness differ with age). A potentially important insight into these determinants comes from a recent study (Heyer et al., 2021) of a sample of epilepsy patients (mean age 39 yrs) who underwent left temporal lobectomy. Histological analysis of resected cortical tissue from the middle temporal gyrus revealed that individual differences in the thickness of this region were driven solely by variation in the thickness of cortical layers II and III. Moreover, increased thickness was associated with fewer, but larger and more highly arborized pyramidal neurons in these layers. Greater thickness was also associated with higher preoperative intelligence scores. These findings offer intriguing clues about the neurobiological underpinnings of the association between cortical thickness and adult cognitive ability. It is currently unknown whether the findings generalize to younger or older populations or to other cortical regions.

Even less is known about the neurobiological correlates of individual differences in GMV, or of brain size more generally. The notion that larger brains confer a cognitive advantage is frequently advanced in an evolutionary context (e.g., Lee et al., 2019; Gonda et al., 2013), but is subject to a number of caveats (e.g., Nave et al., 2018; Striedter, 2005). Not least, much of the variance in human brain volume is associated not with cognitive ability but with sex (see, Deary et al. 2010, for discussion of this apparent paradox). In the present study, for example, despite a GMV that was on average some 10% lower than that of their male counterparts, memory component scores were robustly higher in females (consistent with prior findings, Asperholm et al., 2019; Hirnstein et al., 2022) and no sex difference was evident in mean cognitive ability. Clearly, the factors responsible for sex differences in brain volume are independent of those that mediate its much weaker association with cognitive ability.

In addition to samples of younger and older adults, we also included a small sample of middle-aged individuals. Unsurprisingly, given the modest effect sizes evident even in the older sample for associations between either neural metric and cognitive performance, no significant associations were detected in the middle-aged group. We note however that in regression analyses restricted to the middle-aged and older samples - in which cognitive performance was predicted by sex, years of education, age group, and either cortical thickness or GMV and their interaction with age group – we found no evidence that the strength of the associations between cognitive performance and the neural measures differed significantly between the age groups. Moreover, partial correlations between each measure and cognitive performance differed little from those for the older group alone – for example, for mean cognitive ability, the correlations (controlling for age, sex, and years of education) in the combined group were r = 0.23 (p < 0.002) and r = 0.20 (p < 0.008) for thickness and GMV respectively, as opposed to r = .23 and r = .22 for the older group alone. These findings are consistent with the possibility that the associations with cognitive performance we identified in our older age group extend to middle age. Confirmation of this possibility will of course require the employment of a much larger middle-aged sample than was employed here.

The present study suffers from several limitations. First, the sample sizes are low by contemporary standards, most especially in respect of the middle-aged sample, limiting sensitivity to small effects. Second, the neuropsychological test battery was heavily weighted towards verbal cognition. Notably, there were no tests of spatial or visual memory, and only a single test of reasoning (Progressive Matrices). Third, our study samples were unrepresentative of the general population. Mean cognitive performance was relatively high, and most participants were white, and college educated. Lastly, the study design was cross-sectional, and hence does not allow any inferences to be made about the effects of aging (rather than age) on cognitive or neural metrics and their associations.

These limitations notwithstanding, the present findings add to the existing literature in several ways. Of most significance, the findings indicate that cortical thickness and GMV are non-redundant predictors of cognitive performance, suggesting that the two metrics are neurobiologically distinct correlates of cognitive ability. Furthermore, while confirming a general lack of domain-specificity in associations between cortical thickness and cognitive performance, the findings point to a possible exception in the case of GMV and its association with processing speed. If confirmed, this association deserves further scrutiny. More tentatively, given the post-hoc nature of our analyses, the findings converge with prior research to suggest that the negative associations between cortical thickness and cognitive ability that have consistently been reported in adolescence extend into adulthood, but not beyond about 23-25 years of age.

