## Supplemental Tables for "Cortical thickness, gray matter volume and cognitive performance: a cross-sectional study of the moderating effects of age on their inter-relationships"

Table S1. Hierarchical linear regression results for cortical volume predicting cognitive performance.

| Cortical volume predicting Memory | | | | | | |
| --- | --- | --- | --- | --- | --- | --- |
| Parameter | β | t (p) | F (p) | ΔF (p) | R^2^ | ΔR^2^ |
| Step 1 |  |  | **48.68**  **(< 0.001)** |  | 0.28 |  |
| Intercept |  | **-2.02 (0.045)** |  |  |  |  |
| Age group | **-0.50** | **-10.92**  **(< 0.001)** |  |  |  |  |
| Sex | **-0.26** | **-5.84**  **(< 0.001)** |  |  |  |  |
| Education | **0.12** | **2.59**  **(0.010)** |  |  |  |  |
| Step 2 |  |  | **29.46**  **(< 0.001)** | 0.73 (0.482) | 0.29 | 0.00 |
| Intercept |  | **-2.08 (0.039)** |  |  |  |  |
| Age group | -0.66 | -1.56  (0.120) |  |  |  |  |
| Sex | **-0.29** | **-5.55**  **(< 0.001)** |  |  |  |  |
| Education | **0.12** | **2.64 (0.009)** |  |  |  |  |
| Cortical volume | 0.08 | 1.20 (0.231) |  |  |  |  |
| Cortical volume x age group | 0.21 | 0.49 (0.627) |  |  |  |  |
| Cortical volume predicting Speed | | | | | | |
| Parameter | β | t (p) | F (p) | ΔF (p) | R^2^ | ΔR^2^ |
| Step 1 |  |  | **63.83**  **(< 0.001)** |  | 0.34 |  |
| Intercept |  | **-2.55 (0.011)** |  |  |  |  |
| Age group | **-0.60** | **-13.81**  **(< 0.001)** |  |  |  |  |
| Sex | -0.04 | -0.87  (0.382) |  |  |  |  |
| Education | **0.11** | **2.43**  **(0.016)** |  |  |  |  |
| Step 2 |  |  | **44.92**  **(< 0.001)** | **11.24**  **(< 0.001)** | 0.38 | **0.04** |
| Intercept |  | **-5.21**  **(< 0.001)** |  |  |  |  |
| Age group | **-1.37** | **-3.44**  **(< 0.001)** |  |  |  |  |
| Sex | **-0.14** | **-2.92**  **(0.004)** |  |  |  |  |
| Education | **0.12** | **2.74**  **(0.006)** |  |  |  |  |
| Cortical volume | **0.28** | **4.63**  **(< 0.001)** |  |  |  |  |
| Cortical volume x age group | **0.92** | **2.27**  **(0.024)** |  |  |  |  |
| Cortical volume predicting fluency | | | | | | |
| Parameter | β | t (p) | F (p) | ΔF (p) | R^2^ | ΔR^2^ |
| Step 1 |  |  | **23.17**  **(< 0.001)** |  | 0.16 |  |
| Intercept |  | **-2.55 (0.011)** |  |  |  |  |
| Age group | **-0.41** | **-8.32**  **(< 0.001)** |  |  |  |  |
| Sex | -0.03 | -0.60  (0.552) |  |  |  |  |
| Education | **0.12** | **2.50**  **(0.013)** |  |  |  |  |
| Step 2 |  |  | **15.87**  **(< 0.001)** | **4.30**  **(0.014)** | 0.18 | **0.02** |
| Intercept |  | **-3.86**  **(< 0.001)** |  |  |  |  |
| Age group | -0.61 | -1.34  (0.181) |  |  |  |  |
| Sex | **-0.11** | **-2.00**  **(0.046)** |  |  |  |  |
| Education | **0.14** | **2.73**  **(0.007)** |  |  |  |  |
| Cortical volume | **0.20** | **2.93**  **(0.004)** |  |  |  |  |
| Cortical volume x age group | 0.31 | 0.67  (0.502) |  |  |  |  |
| Cortical volume predicting crystallized IQ | | | | | | |
| Parameter | β | t (p) | F (p) | ΔF (p) | R^2^ | ΔR^2^ |
| Step 1 |  |  | **4.82**  **(0.003)** |  | 0.04 |  |
| Intercept |  | **-2.84 (0.005)** |  |  |  |  |
| Age group | **-0.12** | **-2.35**  **(0.019)** |  |  |  |  |
| Sex | 0.09 | 1.69  (0.092) |  |  |  |  |
| Education | **0.14** | **2.59**  **(0.010)** |  |  |  |  |
| Step 2 |  |  | **3.82**  **(0.002)** | 2.26  (0.105) | 0.05 | 0.01 |
| Intercept |  | **-2.99**  **(0.003)** |  |  |  |  |
| Age group | -0.82 | -1.67  (0.096) |  |  |  |  |
| Sex | 0.04 | 0.69  (0.489) |  |  |  |  |
| Education | **0.14** | **2.62**  **(0.009)** |  |  |  |  |
| Cortical volume | 0.14 | 1.84  (0.067) |  |  |  |  |
| Cortical volume x age group | 0.78 | 1.55  (0.123) |  |  |  |  |
| Cortical volume predicting mean cognitive ability | | | | | | |
| Parameter | β | t (p) | F (p) | ΔF (p) | R^2^ | ΔR^2^ |
| Step 1 |  |  | **41.19**  **(< 0.001)** |  | 0.25 |  |
| Intercept |  | **-2.74 (0.006)** |  |  |  |  |
| Age group | **-0.51** | **-11.05**  **(< 0.001)** |  |  |  |  |
| Sex | **-0.09** | **-1.99**  **(0.048)** |  |  |  |  |
| Education | **0.13** | **2.81**  **(0.005)** |  |  |  |  |
| Step 2 |  |  | **27.09**  **(< 0.001)** | **4.70**  **(0.010)** | 0.27 | **0.02** |
| Intercept |  | **-3.98**  **(< 0.001)** |  |  |  |  |
| Age group | **-1.00** | **-2.32**  **(0.021)** |  |  |  |  |
| Sex | **-0.17** | **-3.14**  **(0.002)** |  |  |  |  |
| Education | **0.14** | **3.00**  **(0.003)** |  |  |  |  |
| Cortical volume | **0.20** | **3.02**  **(0.003)** |  |  |  |  |
| Cortical volume x age group | 0.60 | 1.36  (0.176) |  |  |  |  |

Table S2. Hierarchical linear regression results for white matter volume predicting cognitive performance (results of step 1 models are listed in Table S1).

| White matter volume predicting Memory | | | | | | |
| --- | --- | --- | --- | --- | --- | --- |
| Parameter | β | t (p) | F (p) | ΔF (p) | R^2^ | ΔR^2^ |
| Step 2 |  |  | **29.12**  **(< 0.001)** | 0.12 (0.887) | 0.28 | 0.00 |
| Intercept |  | -1.19 (0.234) |  |  |  |  |
| Age group | -0.33 | -0.91  (0.365) |  |  |  |  |
| Sex | **-0.26** | **-4.92**  **(< 0.001)** |  |  |  |  |
| Education | **0.12** | **2.56 (0.011)** |  |  |  |  |
| White matter volume | -0.01 | -0.15 (0.883) |  |  |  |  |
| White matter volume x age group | -0.17 | -0.48 (0.631) |  |  |  |  |
| White matter volume predicting Speed | | | | | | |
| Parameter | β | t (p) | F (p) | ΔF (p) | R^2^ | ΔR^2^ |
| Step 2 |  |  | **41.38**  **(< 0.001)** | **5.41**  **(0.005)** | 0.36 | **0.02** |
| Intercept |  | **-4.15**  **(< 0.001)** |  |  |  |  |
| Age group | **-0.92** | **-2.70**  **(0.007)** |  |  |  |  |
| Sex | **-0.12** | **-2.37**  **(0.018)** |  |  |  |  |
| Education | **0.10** | **2.32**  **(0.021)** |  |  |  |  |
| White matter volume | **0.16** | **3.23**  **(0.001)** |  |  |  |  |
| White matter volume x age group | 0.33 | 0.98  (0.328) |  |  |  |  |
| White matter volume predicting fluency | | | | | | |
| Parameter | β | t (p) | F (p) | ΔF (p) | R^2^ | ΔR^2^ |
| Step 2 |  |  | **14.60**  **(< 0.001)** | 1.62  (0.198) | 0.17 | 0.00 |
| Intercept |  | **-2.98 (0.003)** |  |  |  |  |
| Age group |  | -0.76  (0.445) |  |  |  |  |
| Sex |  | -1.45  (0.147) |  |  |  |  |
| Education |  | **2.37**  **(0.018)** |  |  |  |  |
| White matter volume |  | 1.74  (0.082) |  |  |  |  |
| White matter volume x age group |  | -0.27  (0.786) |  |  |  |  |
| White matter volume predicting crystallized IQ | | | | | | |
| Parameter | β | t (p) | F (p) | ΔF (p) | R^2^ | ΔR^2^ |
| Step 2 |  |  | **3.01**  **(0.011)** | 0.32  (0.723) | 0.04 | 0.00 |
| Intercept |  | **-2.40**  **(0.017)** |  |  |  |  |
| Age group | -0.31 | -0.76  (0.450) |  |  |  |  |
| Sex | 0.07 | 1.10  (0.270) |  |  |  |  |
| Education | **0.14** | **2.55**  **(0.011)** |  |  |  |  |
| White matter volume | 0.04 | 0.70  (0.484) |  |  |  |  |
| White matter volume x age group | 0.20 | 0.47  (0.638) |  |  |  |  |
| White matter volume predicting mean cognitive ability | | | | | | |
| Parameter | β | t (p) | F (p) | ΔF (p) | R^2^ | ΔR^2^ |
| Step 2 |  |  | **25.20**  **(< 0.001)** | 1.15  (0.317) | 0.26 | 0.00 |
| Intercept |  | **-2.93**  **(0.004)** |  |  |  |  |
| Age group | -0.51 | -1.40  (0.163) |  |  |  |  |
| Sex | **-0.13** | **-2.49**  **(0.013)** |  |  |  |  |
| Education | **0.13** | **2.71**  **(0.007)** |  |  |  |  |
| White matter volume | 0.08 | 1.51  (0.132) |  |  |  |  |
| White matter volume x age group | 0.01 | 0.02  (0.987) |  |  |  |  |

Table S3. Hierarchical linear regression results for total brain volume (adjusted by sex) predicting cognitive performance (results of step 1 models are listed in Table S1).

| Total brain volume predicting Memory | | | | | | |
| --- | --- | --- | --- | --- | --- | --- |
| Parameter | β | t (p) | F (p) | ΔF (p) | R^2^ | ΔR^2^ |
| Step 2 |  |  | **29.36**  **(< 0.001)** | 0.56 (0.572) | 0.29 | 0.00 |
| Intercept |  | **-2.00 (0.046)** |  |  |  |  |
| Age group | **-0.48** | **-9.31**  **(< 0.001)** |  |  |  |  |
| Sex | **-0.26** | **-5.75**  **(< 0.001)** |  |  |  |  |
| Education | **0.12** | **2.63 (0.009)** |  |  |  |  |
| Total brain volume | 0.04 | 0.73 (0.468) |  |  |  |  |
| Total brain volume x age group | 0.04 | 0.89 (0.375) |  |  |  |  |
| Total brain volume predicting Speed | | | | | | |
| Parameter | β | t (p) | F (p) | ΔF (p) | R^2^ | ΔR^2^ |
| Step 2 |  |  | **44.87**  **(< 0.001)** | **11.16**  **(< 0.001)** | 0.38 | **0.04** |
| Intercept |  | **-2.62**  **(0.009)** |  |  |  |  |
| Age group | **-0.50** | **-10.46**  **(< 0.001)** |  |  |  |  |
| Sex | -0.03 | -0.66  (0.507) |  |  |  |  |
| Education | **0.11** | **2.66**  **(0.008)** |  |  |  |  |
| Total brain volume | **0.21** | **4.50**  **(< 0.001)** |  |  |  |  |
| Total brain volume x age group | **0.09** | **2.24**  **(0.026)** |  |  |  |  |
| Total brain volume predicting fluency | | | | | | |
| Parameter | β | t (p) | F (p) | ΔF (p) | R^2^ | ΔR^2^ |
| Step 2 |  |  | **16.05**  **(< 0.001)** | **4.68**  **(0.010)** | 0.18 | **0.02** |
| Intercept |  | **-2.58**  **(0.010)** |  |  |  |  |
| Age group | **-0.33** | **-6.07**  **(< 0.001)** |  |  |  |  |
| Sex | **-0.02** | -0.47  (0.640) |  |  |  |  |
| Education | **0.13** | **2.62**  **(0.009)** |  |  |  |  |
| Total brain volume | **0.16** | **2.97**  **(0.003)** |  |  |  |  |
| Total brain volume x age group | 0.06 | 1.27  (0.205) |  |  |  |  |
| Total brain volume predicting crystallized IQ | | | | | | |
| Parameter | β | t (p) | F (p) | ΔF (p) | R^2^ | ΔR^2^ |
| Step 2 |  |  | **3.67**  **(0.003)** | 1.92  (0.148) | 0.05 | 0.01 |
| Intercept |  | **-2.84 (0.005)** |  |  |  |  |
| Age group | -0.08 | -1.28  (0.200) |  |  |  |  |
| Sex | 0.09 | 1.81  (0.071) |  |  |  |  |
| Education | **0.14** | **2.67**  **(0.008)** |  |  |  |  |
| Total brain volume | 0.09 | 1.63  (0.104) |  |  |  |  |
| Total brain volume x age group | 0.07 | 1.37  (0.171) |  |  |  |  |
| Total brain volume predicting mean cognitive ability | | | | | | |
| Parameter | β | t (p) | F (p) | ΔF (p) | R^2^ | ΔR^2^ |
| Step 2 |  |  | **27.06**  **(< 0.001)** | **4.65**  **(0.010)** | 0.27 | **0.02** |
| Intercept |  | **-2.75**  **(< 0.001)** |  |  |  |  |
| Age group | **-0.44** | **-8.56**  **(< 0.001)** |  |  |  |  |
| Sex | -0.08 | -1.84  (0.066) |  |  |  |  |
| Education | **0.14** | **2.95**  **(0.003)** |  |  |  |  |
| Total brain volume | **0.14** | **2.81**  **(0.005)** |  |  |  |  |
| Total brain volume x age group | 0.08 | 1.69  (0.093) |  |  |  |  |

Results of the linear regression model employing GMV, white matter volume and their interactions to predict speed component scores are shown in Table S4. As is evident from the table, GMV and its interaction with age group significantly predicted speed. By contrast, neither white matter volume nor the white matter volume x age group interaction were predictive of speed.

Table S4. Linear regression results for GMV (adjusted against sex) and white matter volume predicting speed.

| Parameter | β | t (p) | F (p) | R^2^ |
| --- | --- | --- | --- | --- |
|  |  |  | **32.87**  **(< 0.001)** | 0.39 |
| Intercept |  | -1.81  (0.070) |  |  |
| Age group | -0.47 | -1.19  (0.234) |  |  |
| Sex | -0.04 | -0.67  (0.502) |  |  |
| Education | **0.12** | **2.70**  **(0.007)** |  |  |
| GMV | **0.25** | **3.58**  **(< 0.001)** |  |  |
| GMV x age group | **0.11** | **2.29**  **(0.023)** |  |  |
| White matter volume | 0.03 | 0.40 (0.687) |  |  |
| White matter volume x age group | 0.04 | 0.10 (0.918) |  |  |
